# TGF-β induces an atypical EMT to evade immune mechanosurveillance in lung adenocarcinoma dormant metastasis

**DOI:** 10.1101/2024.10.15.618357

**Authors:** Zhenghan Wang, Yassmin Elbanna, Inês Godet, Lila Peters, George Lampe, Yanyan Chen, Joao Xavier, Morgan Huse, Joan Massagué

## Abstract

The heterogeneity of epithelial-to-mesenchymal transition (EMT) programs is manifest in the diverse EMT-like phenotypes occurring during tumor progression. However, little is known about the mechanistic basis and functional role of specific forms of EMT in cancer. Here we address this question in lung adenocarcinoma (LUAD) cells that enter a dormancy period in response to TGF-β upon disseminating to distant sites. LUAD cells with the capacity to enter dormancy are characterized by expression of SOX2 and NKX2-1 primitive progenitor markers. In these cells, TGF-β induces growth inhibition accompanied by a full EMT response that subsequently transitions into an atypical mesenchymal state of round morphology and lacking actin stress fibers. TGF-β induces this transition by driving the expression of the actin-depolymerizing factor gelsolin, which changes a migratory, stress fiber-rich mesenchymal phenotype into a cortical actin-rich, spheroidal state. This transition lowers the biomechanical stiffness of metastatic progenitors, protecting them from killing by mechanosensitive cytotoxic T lymphocytes (CTLs) and natural killer (NK) cells. Inhibiting this actin depolymerization process clears tissues of dormant metastatic cells. Thus, LUAD primitive progenitors undergo an atypical EMT as part of a strategy to evade immune-mediated elimination during dormancy. Our results provide a mechanistic basis and functional role of this atypical EMT response of LUAD metastatic progenitors and further illuminate the role of TGF-β as a crucial driver of immune evasive metastatic dormancy.

## Main

Cancer cells that disseminate from a tumor to distant sites enter a period of dormancy that may persist from months to decades before giving rise to metastasis. Adjuvant therapy treatments seek to prevent metastasis by eliminating residual malignant cells during this dormancy period. Efforts to improve adjuvant therapy are hindered by an insufficient understanding of the molecular mechanisms that preserve the long-term viability of dormant metastatic cells. Identifying these mechanisms is needed to improve treatments and prevent relapse^1-3^.

EMTs are phenotypic plasticity processes that play important roles during development, injury repair, cancer invasion and metastasis^4-6^. During a typical EMT, epithelial cells lose apicobasal polarity, remodel cell adhesion contacts, assemble contractile actin stress fibers, adopt a spindly morphology, gain anteroposterior polarity and become motile. EMT in carcinoma cells is closely associated with migratory and invasive behavior contributing to tumor dissemination and metastasis^7,8^. Despite the current consensus that cancer cells can adopt a spectrum of EMT states with partial or full mesenchymal traits, it remains unclear what determines the extent and purpose of an EMT response during the different phases of metastasis, in particular the dormancy phase.

TGF-β is one of the main drivers of EMT responses in normal and malignant cells^5,9^ and is also an inducer of metastatic dormancy^10-12^. Disseminated carcinoma cells with competence to form dormant metastasis express developmentally primitive progenitor markers, a capacity to enter quiescence, and the ability to elude immune surveillance^13-16^. In experimental models, dormant metastatic cells in brain, lungs, and bone marrow predominantly reside in perivascular niches^17,18^ where TGF-β from the microenvironment induces the entry of these cells into proliferative quiescence^10-12^. In this state, carcinoma cells are thought to evade immune surveillance by downregulating the expression of MHC-I molecules^19-22^, NK cell receptor ligands^13^, and the STING (stimulator of interferon genes) pathway^16^. Reentry of dormant metastatic cells into the cell cycle reactivates the expression of immune activators, leading to cancer cell elimination by CTLs and NK cells^13,16^. These insights suggest that metastatic dormancy is established by malignant progenitors fluctuating between an immune evasive, slow-cycling state and a proliferative state that undergoes immune-mediated elimination^3,23,24^. Importantly, TGF-β plays a critical role in enabling both the invasive phenotype and the immune-evasive dormant state in carcinoma cells, raising questions about the nature of the TGF-β-dependent EMT response accompanying each of these fates.

CTLs and NK cells kill cancer cells by forming an immunological synapse that mediates the delivery of perforin, granzymes, and other pro-apoptotic factors to the target cell^25-28^. Synapse formation depends on the complement of activating ligands on the surface of target cells but is also affected by the biomechanical properties of these cells^29-33^. Increasing the stiffness of target cells through genetic or pharmacological approaches elicits stronger cytotoxic responses, a process termed immune mechanosurveillance^33,34^. In the context of large tumors, TGF-β acts as a direct suppressor of immune effector cells, but during dormancy, solitary disseminated cells persist in the presence of immune surveillance. The biomechanical properties imparted by a classical EMT could render metastatic cells vulnerable to immune mechanosurveillance.

Given the essential roles of TGF-β in metastasis initiation and dormancy^9^, we investigate the EMT state of dormant metastatic cells. Focusing on lung adenocarcinoma (LUAD), a leading cause of cancer mortality globally, our work reveals that dormant metastatic cells undergo a striking morphological transition after extravasation as they undergo TGF-β-induced dormancy. In contrast to the stress fiber-rich, migratory EMT response induced by TGF-β in overtly metastatic cells^35^, we show that dormancy-competent LUAD progenitors respond to TGF-β with an atypical mesenchymal state that is devoid of actin stress fibers. We identify TGF-β-dependent expression of the actin depolymerizing protein gelsolin as a mediator of this transition, and the resulting decrease in cell stiffness allows these cells to avoid mechanosurveillance for the enhanced survival during dormancy. Our results uncovered an atypical EMT response employed by solitary metastatic progenitors to overcome their vulnerability before progressing to overt metastases.

## Results

### A morphological transition in dormant LUAD cells

We previously established the H2087-LCC model of dormant lung adenocarcinoma (LUAD) metastasis by in vivo selection of a latency competent cancer (LCC) cell population derived from a stage I RAS-mutant human LUAD tumor (Extended Data Fig.1a)^13^. When inoculated into the arterial circulation of *Foxn1^nu^* athymic mice, H2087-LCC cells populate perivascular sites in lungs, brain, and other organs, remaining for months as solitary cells or small clusters, and progressing to overt metastasis if the host mice are subjected to NK cell depletion^13^. To generate an analogous immunocompetent model, we applied the LCC selection protocol to 802T4, a cell line derived from a primary LUAD in the *Kras^LSL-G12D/+^;Trp53^flox/flox^* mouse model^36^. The resulting cell population, M802T4-LCC, persisted for months as slow-cycling single cells and small clusters in lungs and brain upon inoculation into immunocompetent B6129SF1/J mice (Extended Data Fig.1b-d). Although recipient mice harboring disseminated M802T4-LCC cells showed few spontaneous outbreaks (Extended Data Fig.1d), antibody-mediated depletion of NK cells, CD4+ or CD8+ T cells in these mice caused widespread metastasis (Extended Data Fig.1e,f). M802T4-LCC cells also formed aggressive metastases when inoculated into immunodeficient NOD scid gamma (NSG) mice (Extended Data Fig.1g). These results validated M802T4-LCC as a model of LUAD metastatic dormancy under immune surveillance. H2087-LCC cells expressed the pluripotency transcription factor (TF) SOX2, which specifies primitive foregut during development^37,38^, and M802T4-LCC cell expressed the proximal lung progenitor TF NKX2-1, which marks early-stage lung bud progenitors (Extended Data Fig.1h)^38,39^. In contrast, cell populations derived from spontaneous metastatic outbreaks in mice (H2087-SO and M802T4-SO cells) expressed the late-stage lung progenitor TF SOX9 predominantly over SOX2 or NKX2-1 (Extended Data Fig.1h), recapitulating a continuum of developmental stages that we previously defined by single-cell transcriptomics in patient-derived LUAD metastasis samples^15^.

To investigate the behavior of dormant H2087-LCC and M802T4-LCC cells in vivo, we analyzed these cells after dissemination to the brain, where the elongated nature of blood capillaries facilitated the morphological analysis of these cells in perivascular niches, and in the lungs, where higher numbers of disseminated cancer cells allowed quantitative analysis. Carcinoma cells extravasate in the brain between 3 days and 7 days after inoculation into the arterial circulation of mice^40,41^. At 7 days after inoculating H2087-LCC into athymic mice and M802T4-LCC into BL/6-albino mice, cancer cells in the brain were situated around blood capillaries and showed a predominantly elongated morphology (Fig.1a,b), as previously observed with other more aggressive models of metastasis^41,42^. This morphology was consistent with reports that circulating cancer cells undergo EMT for migration through endothelia during metastatic extravasation^43^. Notably, the H2087-LCC and M802T4-LCC cells subsequently adopted a spheroidal morphology as they settled into long-term dormancy (Fig.1a,b). The disseminated H2087-LCC and M802T4-LCC cells showed certain mesenchymal traits, including high expression of the mesenchymal marker fibronectin (FN1) and low expression of the epithelial marker E-cadherin, in association with the early elongated morphology as well as the spheroidal morphology that became prevalent weeks after dissemination (Extended Data Fig.2a-c).

**Figure 1.**
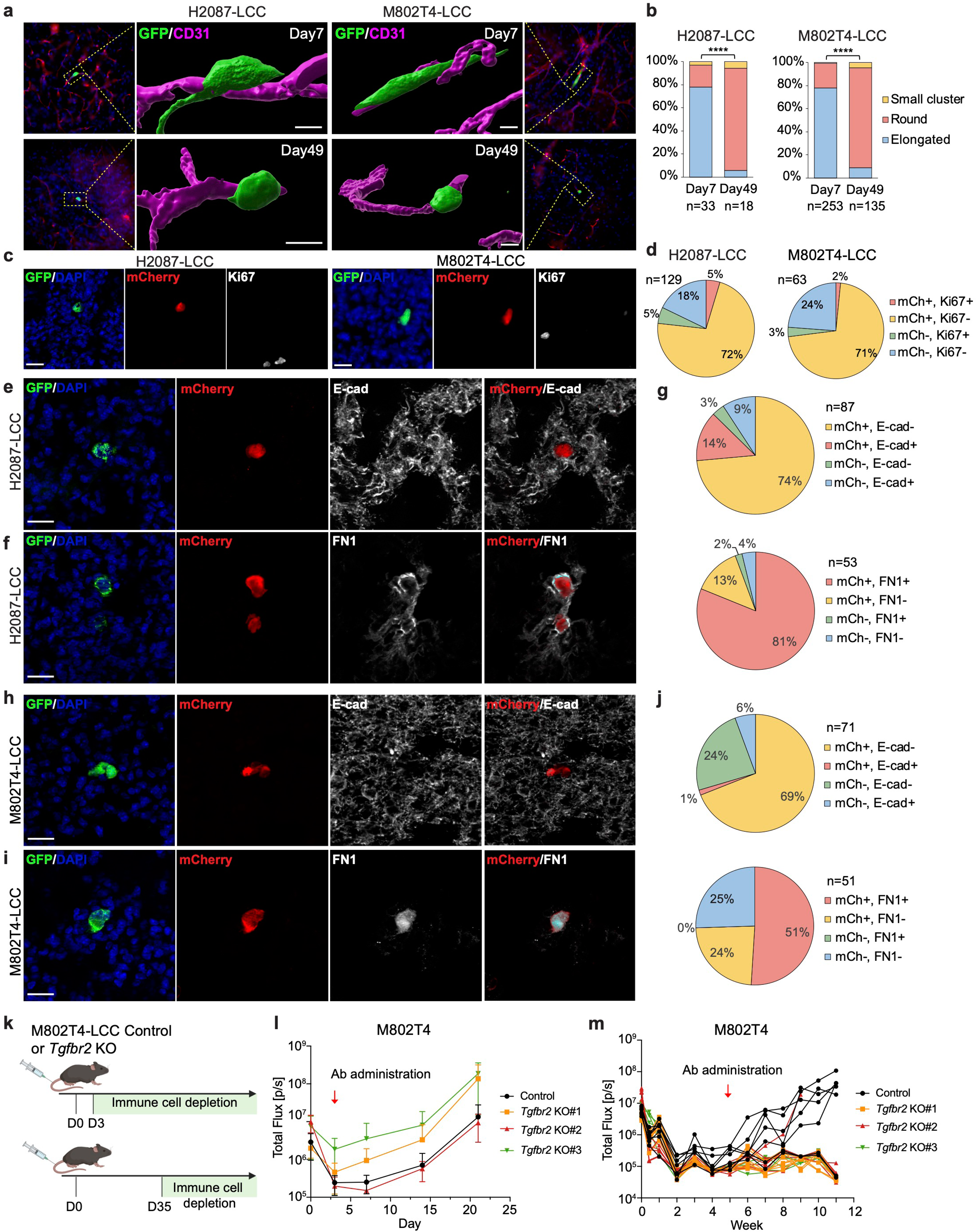
Vital TGF-β signaling in dormant LUAD cells. **a**, 3D reconstructed images of H2087-LCC and M802T4-LCC cells disseminated in brain parenchyma at indicated time point after intracardiac inoculation into athymic mice and B6(Cg)-Tyrc-2J/J (B6-albino) mice, respectively. Cancer cells were visualized by GFP IF (green) and brain capillaries by CD31 IF (magenta). Scale bar: 10 μm. **b**, Quantification of elongated, spheroidal, and microclustered cancer cells (fewer than 10 cells) in A. n=33 and 18 for H2087-LCC cells on day7 and day 49 respectively. n=253 and 135 for M802T4-LCC cells on day 7 and day 49 respectively. Chi-square test. **c**, Representative images of H2087-LCC and M802T4-LCC cells expressing a TGF-β reporter in the lungs of athymic mice and B6129SF1/J mice, respectively, 7 weeks after intravenous injection. Cancer cells were visualized by GFP IF (green), the TGF-β mCherry reporter activity in red, Ki67 proliferation marker in white, and DAPI in blue. Scale bar: 20 μm. **d**, Quantification of TGF-β mCherry reporter and Ki67 expression in H2087-LCC (n=129) and M802T4-LCC (n=63) cells in the lungs of athymic and B6129SF1/J mice, respectively, 7 weeks after intravenous injection. **e** and **f**, Representative IF images of H2087-LCC in the lungs of athymic mice 7 weeks after intravenous injection. GFP^+^ cancer cells (green), TGF-β mCherry reporter (red), E-cadherin or FN1 (white), and DAPI (blue). **g**, Quantification of TGF-β mCherry reporter and E-cadherin (n=87) or FN1 (n=53) expression in disseminated H2087-LCC cells in the lungs of athymic mice. **h** and **i**, Representative IF images of M802T4-LCC in the lungs of B6129SF1/J mice 7 weeks after intravenous injection. GFP^+^ cancer cells (green), TGF-β mCherry reporter (red), E-cadherin (white), and DAPI (blue). Scale bar: 20 μm. **j**, Quantification of TGF-β mCherry reporter and E-cadherin (n=71) or FN1 (n=51) expression in M802T4-LCC in the lungs of B6129SF1/J mice 7 weeks after intravenous injection. **k-m**, Schematic representation of the experimental design, created with Biorender.com (k), and tracking of wild-type and *Tgfbr2* KO M802T4-LCC cells intravenously inoculated in B6129SF1/J mice followed by antibody-mediated depletion of NK, CD4+ and CD8+ T cells from day 3 (l) or day 35 (m) after inoculation. n=6 or 7 mice per group.

### Vital TGF-β signaling in dormant LUAD cells

TGF-β produced by endothelial and perivascular cells is known to mediate the entry of disseminated carcinoma cells into a dormant state^23^. To determine if dormant solitary LUAD cells persistently harbored TGF-β signaling activity, we engineered a doxycycline-dependent TGF-β mCherry reporter construct^16^ into H2087-LCC and M802T4-LCC cells (Extended Data Fig.2d,e). Mice were inoculated with these cells and switched to doxycycline-containing chow to allow TGF-β reporter expression. Long-term disseminated H2087-LCC and M802T4-LCC in the lungs (Fig.1c,d) and brain (Extended Data Fig.2f,g) showed persistent mCherry expression together with low expression of the cell proliferation marker Ki67, as well as low expression of E-cadherin and high expression of FN1 (Fig.1c-j). These results indicate that long-term dormant LUAD cells harbored TGF-β signaling activity associated with cell quiescence and mesenchymal traits during prolonged dormancy period.

To evaluate the importance of TGF-β signaling for LUAD dormancy, we intravenously inoculated B6129SF1/J immunocompetent mice with wild-type M802T4-LCC cells or M802T4-LCC cells with CRISPR-Cas9 knockout of the TGF-β receptor *Tgfbr2* (Extended Data Fig.2h,i). We then subjected the mice to antibody-mediated depletion of NK cells, CD4^+^ T, and CD8^+^ T cells either 3 days or 35 days after inoculation, to allow the metastatic outgrowth of disseminated cancer cells (Fig.1k). This protocol provides a readout of metastasis-initiating competence remaining in latent cancer cell populations after specific perturbations^13,16^. Depletion of NK and T cells early (day 3) after cancer cell inoculation led to the aggressive outgrowth of metastatic colonies in the lungs of all the inoculated mice, which was not inhibited by the knockout of *Tgfbr2* (Fig.1l). This result indicates that TGF-β signaling was not essential for metastatic seeding of M802T4-LCC cells in the lungs. Depletion of NK and T cells 35 days after inoculation led to metastatic outbreaks in 5 out of 7 mice harboring disseminated M802T4-LCC cells. However, this late depletion led to metastatic outbreaks in only 1 out of 21 mice harboring disseminated *Tgfbr2*-KO M802T4-LCC cells (*p* = 0.0012, Fisher’s exact test) (Fig.1m). These results suggest that TGF-β is important for preservation of metastasis-initiating M802T4-LCC cells during prolonged dormancy.

### An EMT state lacking actin stress fibers

TGF-β is a potent inducer of EMTs in normal and malignant epithelial cells^9^. Aggressive metastatic cells derived from LUAD patients and KP mouse models respond to TGF-β by undergoing a typical EMT coupled with expression of a set of fibrogenic factors, which is critical for the ensuing metastatic outgrowth^35^. Although the spindly morphology of H2087-LCC and M802T4-LCC cells upon extravasation (refer to Fig.1a) was compatible with a classical EMT, the spheroidal configuration adopted by these cells during long-term dormancy was not consistent with a typical EMT. Mesenchymal traits without an overt EMT state have also been noted in dormant breast cancer cells^44-46^. These observations raised questions about the fate of the EMT response in dormancy-competent LUAD cells under sustained TGF-β stimulation.

To investigate this question, we incubated H2087-LCC and M802T4-LCC cells with TGF-β (100 pM TGF-β1) or the TGF-β receptor inhibitor SB505124^47^ which sets a baseline by inhibiting endogenous TGF-β signaling in the cultures. Both cell models responded to TGF-β by initially adopting a spindly morphology (Fig.2a,b and Extended Data Fig.3a,b) accompanied with a loss of E-cadherin, β-catenin, and ZO-1 expression and a gain of actin stress fibers (Fig.2c and Extended Data Fig.3c) and motility (Extended Data Fig.3d and Video 1), all hallmarks of a typical full EMT^6^. Notably, after 3 days of incubation with TGF-β, the LCC cells started to transition into a spheroidal morphology lacking stress fibers (Fig. 2a,c and Extended Data Fig.3a,b) and motility (Extended Data Fig. 3d,e and Video 1), a phenotype that became prevalent after 7 days of incubation with TGF-β. The actin stress fibers appearing during the early phase of the TGF-β response were replaced with subcortical actin filaments by day 7, consistent with the observed transition from an elongated and motile to a spheroidal and steady morphology (Fig. 2c and Extended Data Fig.3c). This morphological transition resembled that of H2087-LCC and M802T4-LCC cells entering dormancy in vivo. In contrast, H2087-SO and M802T4-SO derivatives exhibited typical EMT traits during prolonged incubation with TGF-β (Fig. 2a,b and Extended Data Fig. 3a,b,f,g) similar to those of aggressively metastatic LUAD cell lines^35^.

**Figure 2.**
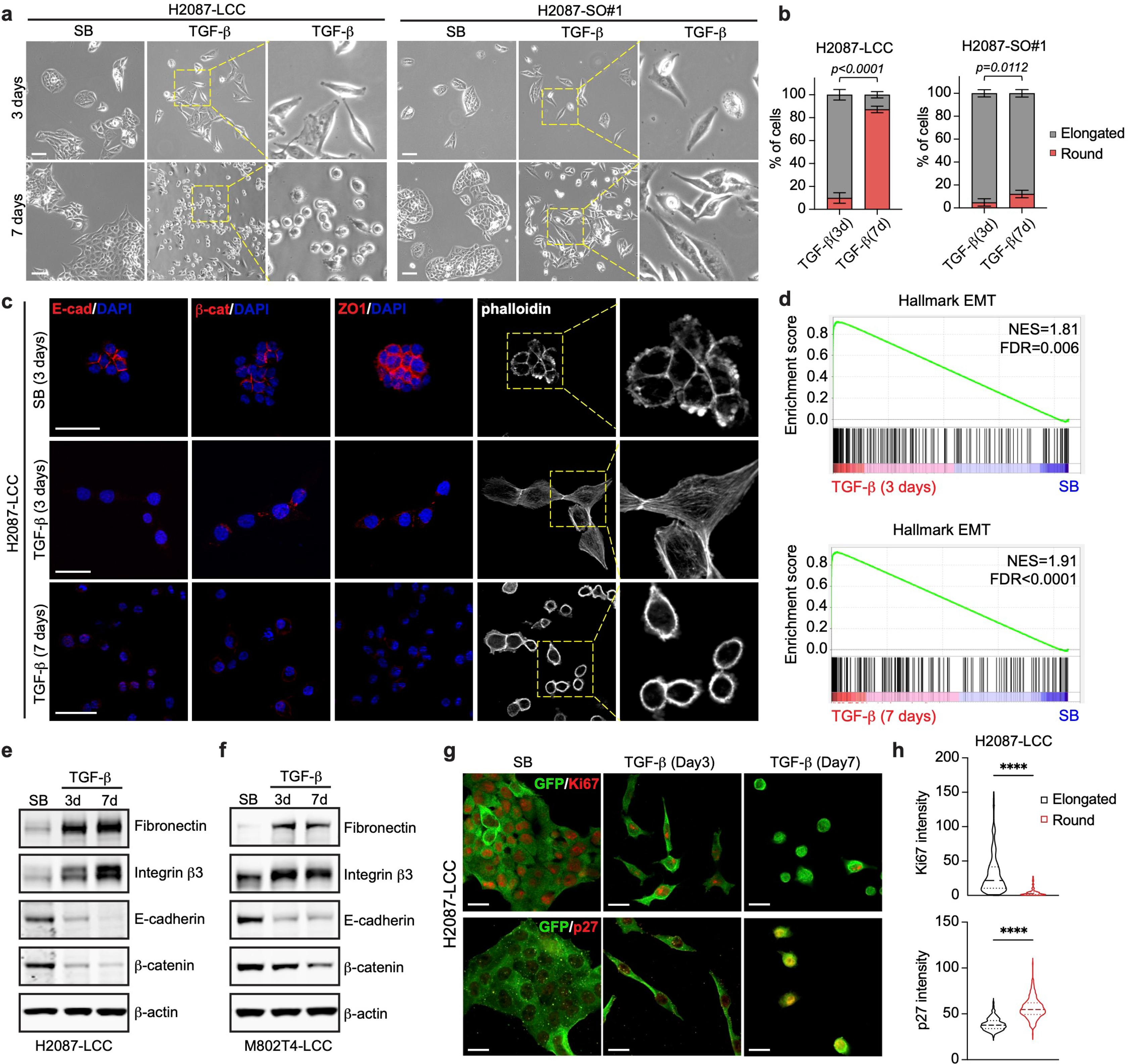
**An EMT state lacking actin stress fibers**. **a**, Representative bright-field images of H2087-LCC and H2087-SO cell cultures after incubation with TGF-β for 3 days and 7 days. Scale bar: 200 μm. **b**, Quantification of elongated and spheroidal cells in A. Cells from 10 images were analyzed for each condition. Data are mean ± SD.; two-sided unpaired *t*-test. **c**, Representative IF images for the epithelial markers E-cadherin, β-catenin, and zona occludens 1 (ZO-1), and of phalloidin staining in H2087-LCC cells incubated with TGF-β for 3 days or 7 days. Scale bar: 20 μm. **d**, Expression of a hallmark EMT gene signature in H2087-LCC cells after incubation with TGF-β for 3 days and 7 days. **e** and **f**, Western immunoblot analysis of the indicated proteins in H2087-LCC and M802T4-LCC cells after incubation with TGF-β for 3 days or 7 days, or with SB for 3 days. β-actin was used as loading control. **g**, Representative IF images of H2087-LCC cells after incubation with TGF-β for 3 days or 7 days. GFP^+^ cancer cells (green), Ki67 (red) or p27KIP1 (red). Scale bar: 20 μm. **h**, Quantification of Ki67 and p27 IF intensity in elongated H2087-LCC cells after 3 days of incubation with TGF-β and spheroidal H2087-LCC cells after 7 days of incubation with TGF-β. n=299 (elongated) and n=116 (round) for Ki67 staining, n=222 (elongated) and n=265 (round) for p27 staining. Two-sided unpaired *t*-test.

### Distinct EMT responses in different LUAD progenitor states

Despite the lack of stress fibers and motility, spheroidal H2087-LCC and M802T4-LCC cells continued to show low levels of E-cadherin, β-catenin, and ZO-1 IF staining after 7 days of incubation with TGF-β (Fig. 2c and Extended Data Fig. 3c). Gene set enrichment analysis of RNA sequencing (RNA-seq) data demonstrated the persistence of a strong EMT transcriptional program in H2087-LCC cells incubated with TGF-β for 7 days (Fig. 2d). Analysis confirmed an increase in fibronectin and integrin β3 (ITGB3) expression as mesenchymal state markers, and a downregulation of E-cadherin and β-catenin expression in H2087-LCC and M802T4-LCC cells incubated for 3 days or 7 days with TGF-β (Fig. 2e,f). The transcriptomic profiling of H2087-LCC cells incubated with TGF-β for 7 days also revealed an inhibition of biosynthetic processes and DNA replication compared with cells treated with TGF-β for 3 days (Table 1). Entry of these cells into a slow-cycling state was confirmed by a low expression of the proliferation marker Ki67 and a high expression of the quiescence marker p27KIP1^48^ in LCC cells upon transitioning into the spheroidal shape in response to TGF-β (Fig. 2g,h and Extended Data Fig. 4a,b). H2087-SO cells showed a less pronounced growth inhibitory response (Extended Data Fig.4c). H2087-LCC cells placed in regular media after a 7-day incubation with TGF-β transitioned back to an epithelial morphology, indicating reversibility of the effect, whereas cells that continued with TGF-β for up to 14 days retained a spheroidal morphology (Extended Data Fig.4d).

We previously showed that TGF-β signaling blunts cell proliferation partly through induction of CDK inhibitors^49,50^. TGF-β treatment increased the expression of *CDKN1A* (encoding p21CIP1) and *CDKN2B* (encoding p15INK4B) inhibitors in H2087-LCC cells (Extended Data Fig.4e). We also showed that under growth restrictive conditions, H2087-LCC cells additionally express the WNT inhibitor DKK1 which mediates resistance to WNT stimulation and enforces dormancy in vivo^13^. We found that incubation with TGF-β strongly increased DKK1 mRNA and protein expression in H2087-LCC cells but not in H2087-SO cells (Extended Data Fig.4f,g). H2087-SO cell populations showed a diminished ability to express DKK1 and resist WNT3A induction of the canonical WNT target gene *Axin2* compared to H2087-LCC cells (Extended Data Fig.4h).

In aggressive metastatic cells from LUAD and other types of carcinoma, TGF-β induces the expression of the EMT TF Snail (encoded by *Snai1*) coupled with the expression of a set of fibrogenic factors including interleukin-11 (*Il11*), hyaluronan synthase 2 (*Has2*), serpin E1 (*Serpine1*), and others^35,51^. Both the EMT and the fibrogenic arms of this response support the metastatic outgrowth of aggressive LUAD cells in the lungs^35,52^. These EMT-associated fibrogenic gene responses were present, albeit attenuated in H2087-LCC and M802T4-LCC cells compared to these responses in the highly metastatic cell lines A549, derived from a KRAS-mutant human LUAD^53^, and 393T3, derived from an aggressive KP mouse LUAD tumor (Extended Data Fig. 5a,b)^36^. The fibrogenic EMT response to TGF-β in aggressive LUAD cancer cells depends on RAS/MAPK signaling through RREB1 (Ras response element binding protein 1). RREB1 directly cooperates with TGF-β-activated SMAD TFs to stimulate primed enhancers in the EMT-TF and fibrogenic target genes^35,52^. A similar RREB1 dependence of these TGF-β gene responses in H2087-LCC cells was observed upon knock down of *RREB1* (Extended Data Fig.5c,d).

In sum, H2087-LCC and M802T4-LCC populations rich in SOX2^+^ and NKX2-1^+^ primitive LUAD progenitors respond to TGF-β with deployment of an atypical state characterized by a robust EMT transcriptional program, a spheroidal morphology lacking actin stress fibers and motility, an attenuated set of fibrogenic gene responses, and proliferative quiescence. This contrasts with the persistent induction of a typical EMT with a strong fibrogenic response by TGF-β in developmentally more advanced LUAD cells, showing that TGF-β triggers distinct EMT responses in primitive (SOX2^+^/NKX2-1^+^) versus late-stage (SOX9^+^) LUAD progenitors.

### TGF-β increases gelsolin expression in LUAD progenitors

To investigate the basis for the loss of actin stress fibers in LCC cells under persistent exposure to TGF-β, we queried our RNA-seq dataset for differentially expressed actin cytoskeleton components and regulators. *GSN* (gelsolin), *MYLK2* (myosin light chain kinase 2) and *ITGB3* emerged as differentially expressed genes in H2087-LCC cells incubated with TGF-β for 7 days, when most cells had transitioned into the spheroidal morphology, compared with cells incubated with TGF-β for 3 days, when most cells were still spindly (Fig.3a). In agreement with these results, H2087-LCC and M802T4-LCC cells incubated with TGF-β showed a progressive increase in gelsolin levels which was only incipient after 3 days and reached 2-to 3-fold over basal levels after 7 days both at the protein (Fig.3b) and mRNA levels (Fig.3c). In contrast, the protein levels of MYLK2 (Extended Data Fig.6a) and ITGB3 (Fig. 2e,f) showed little or no further increase after 7 days versus 3 days of incubation with TGF-β. Notably, TGF-β did not increase *GSN* expression in SO derivatives or in the aggressive metastatic LUAD cell lines A549 or 393T3 (Fig.3c).

**Figure 3.**
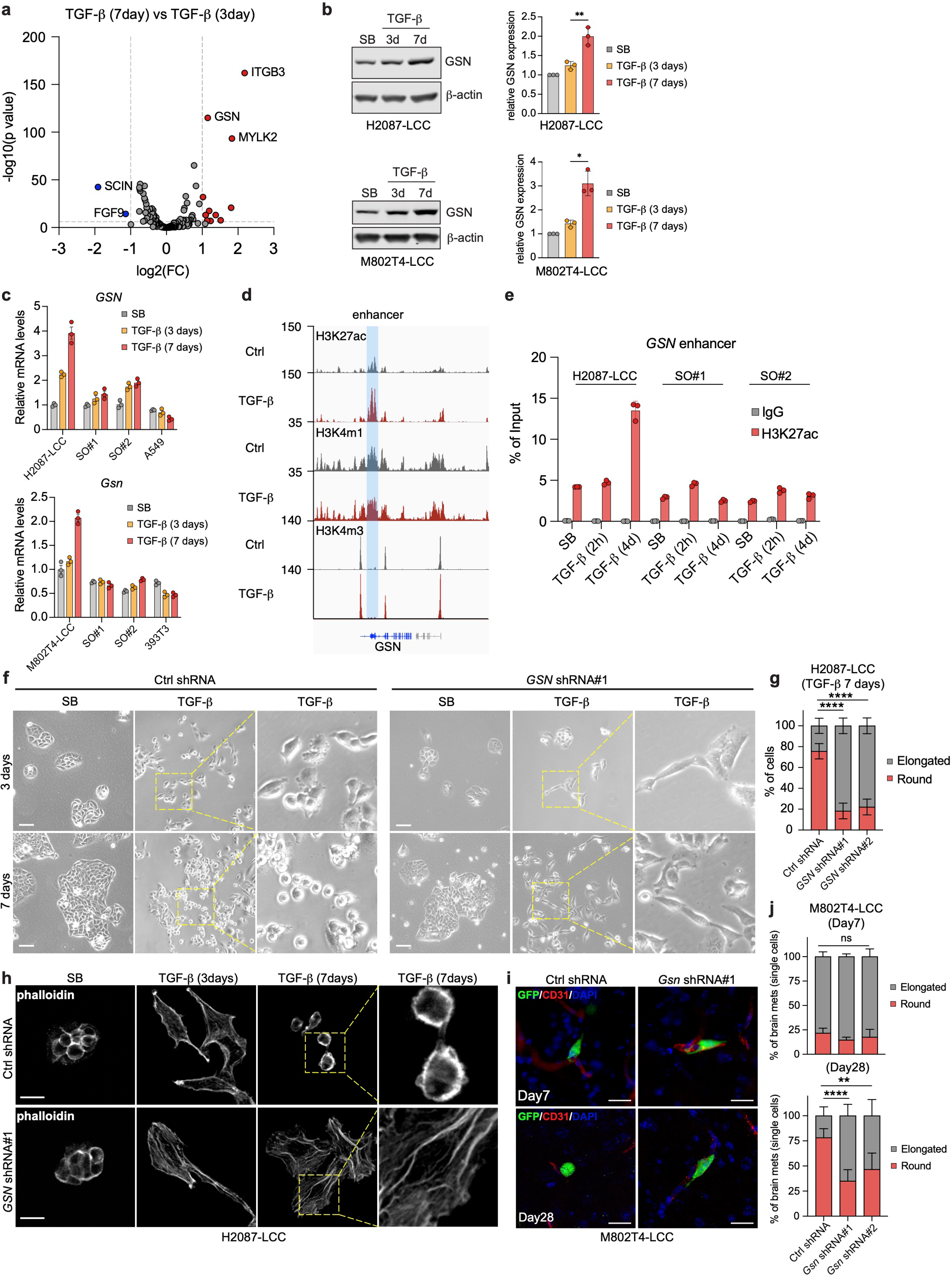
A gelsolin modified EMT. **a**, Volcano plot of differentially expressed actin cytoskeleton components and regulators in H2087-LCC cells incubated with TGF-β for 7 days versus 3 days. **b**, Western immunoblot analysis (left) and quantification (right) of gelsolin levels in H2087-LCC and M802T4-LCC cells treated with TGF-β for the indicated time periods. β-actin was used as loading control. Quantification was done with three independent experiments. Data are mean ±SD., two-sided unpaired *t*-test. **c**, qRT-PCR analysis of *GSN* mRNA levels in the indicated cell lines and TGF-β treatment conditions. Data are mean ± SEM; n=3. **d**, Gene track views of H3K27ac, H3K4m1 and H3K4m3 ChIP-seq tags at the *GSN* locus in H2087-LCC cells incubated with TGF-β for 4 days. The shared region shows histones marks typical of an enhancer. **e**, ChIP-PCR analysis of H3K27ac at the *GSN* enhancer in the indicated cell lines upon incubations with SB or TGF-β for 2h or 4 days. Data are mean ± SEM; n=3. **f**, Representative bright-field images of H2087-LCC cells expressing control shRNA or *shGSN* and incubated with TGF-β for 3 days or 7 days. Scale bar: 200 μm. **g**, Quantification of elongated versus spheroidal H2087-LCC cells in F. Cells from 10-12 images were analyzed for each condition. Data are mean ± SD.; two-sided unpaired *t*-test. **h**, Representative phalloidin staining of H20878-LCC cells expressing control shRNA or *shGSN* and incubated with TGF-β for 3 days or 7 days. Scale bar: 20 μm. **i**, Representative IF images of M802T4-LCC cells expressing the indicated shRNAs, in the brain of Albino-B6 mice 7 days or 28 days after intracardiac inoculation. GFP^+^ cancer cells (green), CD31^+^ capillaries (red), DAPI (blue). Scale bar: 20 μm. **j**, Quantification of elongated versus spheroidal M802T4-LCC cells in the brain parenchyma of Albino-B6 mice that were intracardially inoculated with these cells 7 days or 28 days prior. Results from n=4 mice (Day7) or n=5 mice (Day28) per group. Chi-square test.

We previously profiled histone 3 (H3) modifications in H2087-LCC cells using chromatin immunoprecipitation with sequencing (ChIP-seq)^16^. H3K4me3, which marks active promoters^54,55^, was enriched near the *GSN* transcription start site independently of TGF-β treatment. H3K4me1 and H3K27ac, which mark active enhancers, were present in a *GSN* intronic region prior to TGF-β addition. Incubation with TGF-β for 4 days increased the level of H3K27ac in this region (Fig.3d). ChIP-PCR analysis showed that TGF-β induced an increase in H3K27 acetylation level in this enhancer region in H2087-LCC cells, but not in H2087-SO cells derived from spontaneous metastatic outbreaks (Fig.3e). Collectively, these results suggest that TGF-β induces a progressive and specific increase in gelsolin expression in latency competent cells.

### A gelsolin-modified EMT

Gelsolin promotes actin cytoskeleton turnover through severing and capping of actin filaments^56,57^. The observed loss of actin stress fibers in cells incubated with TGF-β long-term was consistent with a role of gelsolin as a mediator of this transition. To investigate this, we knocked down gelsolin expression with two different shRNAs in H2087-LCC and M802T4-LCC cells (Extended Data Fig.6b), which did not inhibit the induction of *Snai1* by TGF-β (Extended Data Fig.6c). The depletion of gelsolin did not interfere with the initial induction of actin stress fibers or a spindly cell morphology by TGF-β, but it consistently prevented the transition from spindly to spheroidal morphology after prolonged incubation with TGF-β (Fig.3f,g and Extended Data Fig.6d,e). Phalloidin staining of actin filaments confirmed that gelsolin knockdown inhibited the TGF-β-induced transition of actin stress fibers to cortical actin filaments (Fig.3h). Unlike the gelsolin knockdown, knockdown of ITGB3 or MYLK2 did not affect the transition of H2087-LCC cell morphology from spindly to spheroidal during incubation with TGF-β (Extended Data Fig. 6f,g).

To determine the effect of gelsolin depletion on the ability of disseminated cancer cells to transition to a spheroidal morphology in vivo, we performed quantitative morphometry in M802T4-LCC cells disseminate to the brain. The knockdown of gelsolin had no effect on the predominantly elongated morphology of recently extravasated cells compared to controls one week after intracardiac inoculation of cells into mice (Fig.3i,j). However, 28 days after inoculation, the predominantly spheroidal morphology of isolated M802T4-LCC cells in the brain was significantly reduced in gelsolin knockdown cells compared to control cells (Fig. 3i,j). Collectively, these results identify gelsolin as a critical mediator of the morphological transition of dormancy-competent LUAD cells from an elongated, stress fiber-rich mesenchymal state during the early phase of a TGF-β response to a cortical actin-rich mesenchymal state after a prolonged exposure to TGF-β in vitro and in vivo.

### An immune evasive decrease in cell stiffness

The state of the cytoskeleton determines the stiffness of a cell^58^, and a high level of stiffness in cancer cells favors the formation of the cytotoxic synapse and killing by CTLs and NK cells^28,31-33,59^. The dramatic switch from a stress fiber-rich state to a spheroidal, cortical actin-rich state induced by TGF-β in dormant metastatic cells raised the possibility that this transition regulates the susceptibility of the cells to immune mechanosurveillance during long-term dormancy.

We used atomic force microscopy (AFM)^60^ to measure surface tension and determine the stiffness of LUAD cells under different TGF-β treatment conditions in culture (Fig.4a). LCC cells gained stiffness as they adopted a spindly morphology upon incubation with TGF-β for 3 days. The level of cell stiffness then declined as the cells adopted a spheroidal morphology under continued incubation with TGF-β (Fig.4b,c). To determine if this transition to a soft morphology was associated with a change in cell susceptibility to immune-mediated killing, we treated H2087-LCC and M802T4-LCC cells with TGF-β and co-cultured the cells with interleukin 2 (IL-2)-activated mouse NK cells (Fig.4d). Cells incubated with TGF-β for 3 days were more susceptible to NK cell-mediated killing than were cells incubated for 7 days (Fig.4e). Similar results were obtained when H2087-LCC cells were co-cultured with human NK cells isolated from human peripheral blood from different donors (Fig.4f). Live imaging of H2087-LCC cell cultures incubated with TGF-β for 5 days, which included a mix of spindly and spheroidal cells, revealed that NK cells interacted with both cell morphologies but were more effective at killing the spindly cancer cells (Fig.4g and Video 2).

**Figure 4.**
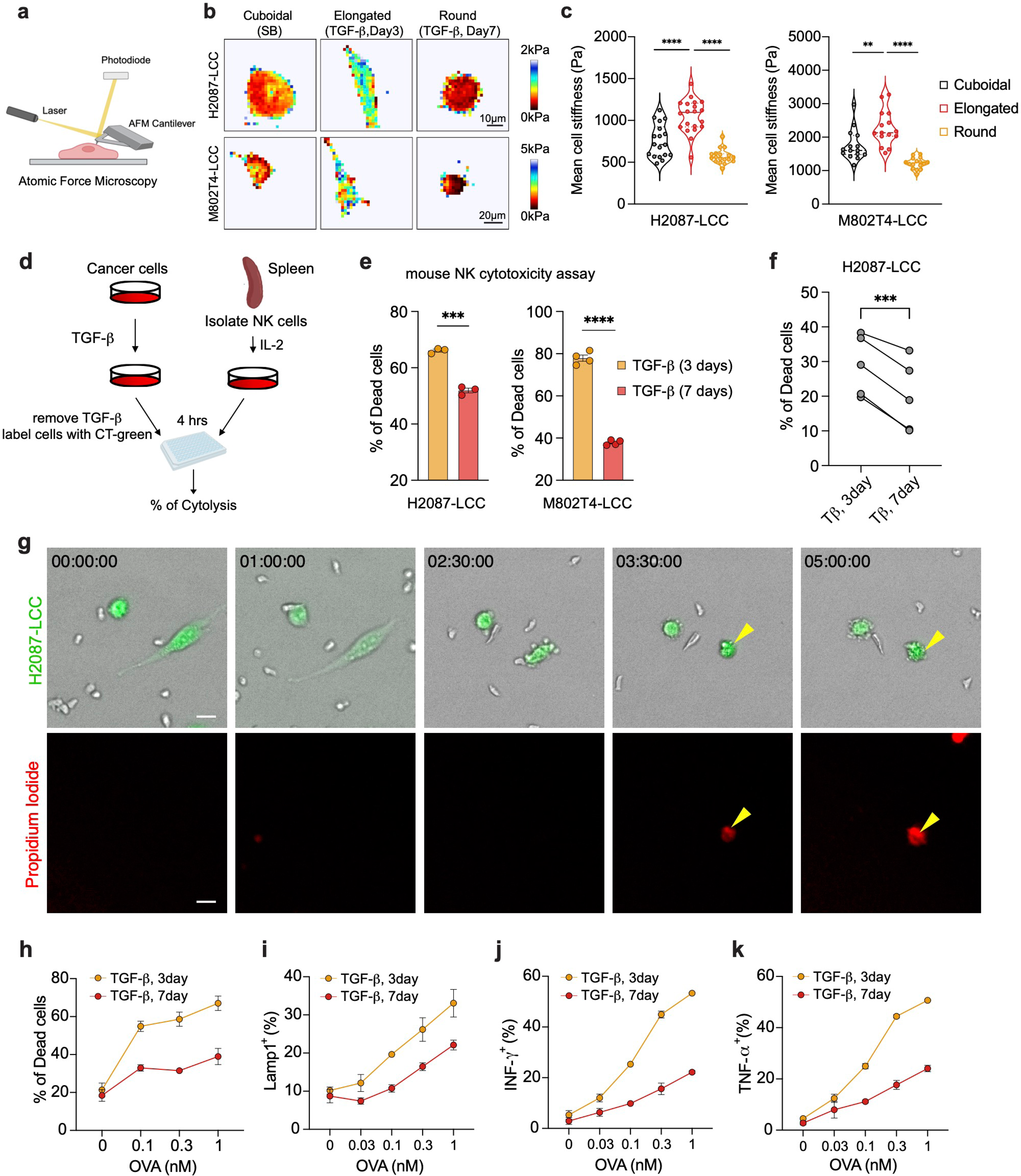
An immune evasive decrease in cell stiffness. **a**, Schematic representation of atomic force microscopy (AFM) determination of the biomechanical stiffness of cells (created with Biorender.com). **b**, Force maps of representative cells, with Young’s modulus value (kPa) indicated in pseudocolor. **c**, Mean cell stiffness of elongated and spheroidal H2087-LCC or M802T4-LCC cells in response to TGF-β treatment for 3 days and 7 days. n=18 per condition for H2087-LCC cells; n=15 per condition for M802T4-LCC cells. Two-sided unpaired *t*-test. **d**, Schematic of in vitro mouse NK cell cytotoxicity assay (created with Biorender.com). **e**, H2087-LCC and M802T4-LCC cells were incubated with TGF-β for the indicated time periods and admixed with IL-2-activated mouse NK cells. Data are mean ± SEM, n=3 or 4; two-sided unpaired *t*-test. **f**, H2087-LCC cells were incubated with TGF-β for the indicated time periods and admixed with NK cells derived from human peripheral blood from 5 different donors. After 4 h of coincubation, the percentage of dead cancer cells were determined by flow cytometry. Connecting lines indicate samples derived from the same donor. Paired *t*-test. **g**, H2087-LCC cells were incubated with TGF-β for 5 days to generate a mixed of cells with elongated or spheroidal morphology. The cells were then admixed with IL-2-activated mouse NK cells in the presence of propidium iodide, which stains the nuclear DNA of dead cells. Shown are still frames from time-lapse imaging of H2087-LCC and NK cell cocultures. Arrowhead, an elongated cancer cell undergoing death near a spheroidal cancer cells that survives NK attack. Scale bar: 50 μm. **h**, M802T4-LCC cells incubated with TGF-β for 3 days or 7 days were loaded with increasing concentrations of OVA peptide and then mixed with OT1 CTLs. Cancer cell lysis was quantified after 5 h. Data are mean ± SD; n=3. **i**, CTL degranulation as determined by surface exposure of Lamp1 5 h after mixing OT1 CTLs with OVA-loaded M802T4-LCC cells. **j** and **k**, Production of IFN-γ and TNF-α, measured by intracellular immunostaining of CTLs 5 h after mixing with the indicated M802T4-LCC cells. Data are mean ± SD; n=3.

To determine whether the TGF-β dependent change in cell morphology was associated with a difference in killing by CTLs, we incubated M802T4-LCC cells with TGF-β for 3 days or 7 days before adding ovalbumin_257-264_ peptide (OVA), which binds to the class I MHC molecule H-2K^b^ and renders cells susceptible to killing by CTLs expressing the OVA T-cell receptor OT1^61^. When co-cultured with OT1 CTLs, cancer cells were more susceptible to killing after incubation with TGF-β for 3 days than for 7 days (Fig.4h). Moreover, cells incubated with TGF-β for 3 days induced stronger CTL activation, as indicated by increased degranulation (measured by the lysosomal marker Lamp1) (Fig.4i) and increased production of the inflammatory cytokines interferon-*γ* and tumor necrosis factor (TNF) (Fig.4j,k).

The ADP-ribosyl transferase domain (a.a. 375-591; referred to as DeAct) of *Salmonella enterica* SpvB drives the disassembly of filamentous actin by catalyzing the ADP-ribosylation of actin on Arg177^62^. To establish causality between actin cytoskeleton depolymerization and evasion of immune surveillance, we engineered H2087-LCC and M802T4-LCC cells with doxycycline-inducible DeAct expression as a genetically encoded actin disassembly tool (Fig.5a). During the early phase (day 3) of cell incubation with TGF-β, cells expressing DeAct (marked by mCherry) for 24 h showed a lack of actin stress fibers relative to DeAct negative controls (Fig.5b). Importantly, DeAct-expressing cells exhibited a dramatic reduction in cell stiffness (Fig.5c) and were less sensitive to both NK cell-mediated and CTL-mediated killing compared with control cells expressing the empty vector (Fig.5d,e). These results indicated that actin cytoskeleton-dependent cell stiffness regulates mechanosurveillance in these LUAD cells.

**Figure 5.**
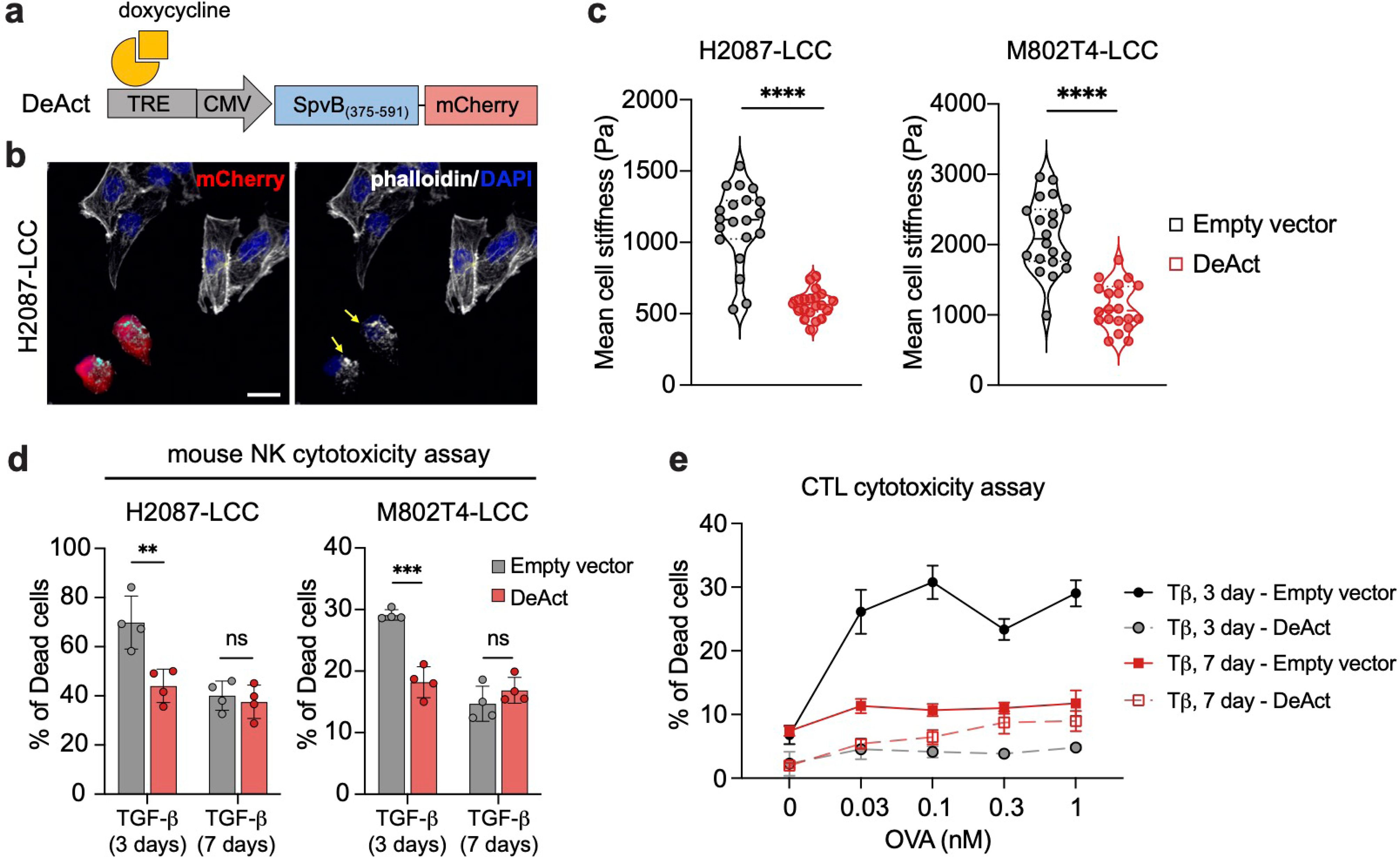
Actin depolymerization lowers cancer cell stiffness and immune-mediated killing. **a**, Schematic representation of a vector driving Dox-inducible co-expression of the DeAct SpvB_375-591_ and a mCherry marker. **b**, Representative images of H2087-LCC cells incubated with doxycycline for 24 h to induce DeAct and mCherry expression in transduced H2087-LCC cells admixed with non-transduced cells. Phalloidin staining demonstrates absence of actin stress fibers in DeACt expressing cells. **c**, Mean cell stiffness in H2087-LCC and M802T4 cells incubated with TGF-β for 3 days to induce an EMT and express control vector or DeAct for 24h before analysis. n=20 per condition; two-sided unpaired *t*-test. **d**, cytotoxic effect of mouse NK cell on the indicated cancer cells incubated with TGF-β for 3 days or 7 days and express control vector or DeAct 24h before analysis. Data are mean ± SEM, n=4; two-sided unpaired *t*-test. **e**. cytotoxic effect of CTLs on M802T4-LCC cells incubated with TGF-β for 3 days or 7 days and express control vector or DeAct 24h before analysis. Data are mean ± SD; n=3.

### Gelsolin protects dormant metastasis from mechanosurveillance

Next, we tested the hypothesis that TGF-β allows dormant progenitor cells to evade mechanosurveillance through a gelsolin-mediated transition from a typically mesenchymal spindly morphology into a spheroidal morphology. AFM measurements showed that knockdown of gelsolin caused the retention of a high level of stiffness in cells incubated with TGF-β for 7 days, both in the H2087-LCC (Fig.6a,b) and M802T4-LCC models (Extended Data Fig.7a,b). The knockdown of gelsolin rendered these cells more vulnerable to NK cell mediated killing (Fig.6c and Extended Data Fig.7c). Similarly, knocking down gelsolin expression in M802T4-LCC cells drove stronger degranulation of CTLs and cytokine production (Extended Data Fig.7d-f), and resulted in increased killing of cancer cells in co-culture with CTLs (Extended Data Fig.7g). Importantly, the expression of DeAct reversed the effect of gelsolin knockdown and rescued NK cell-mediated killing in both H2087-LCC and M802T4-LCC cells (Fig.6d-g).

**Figure 6.**
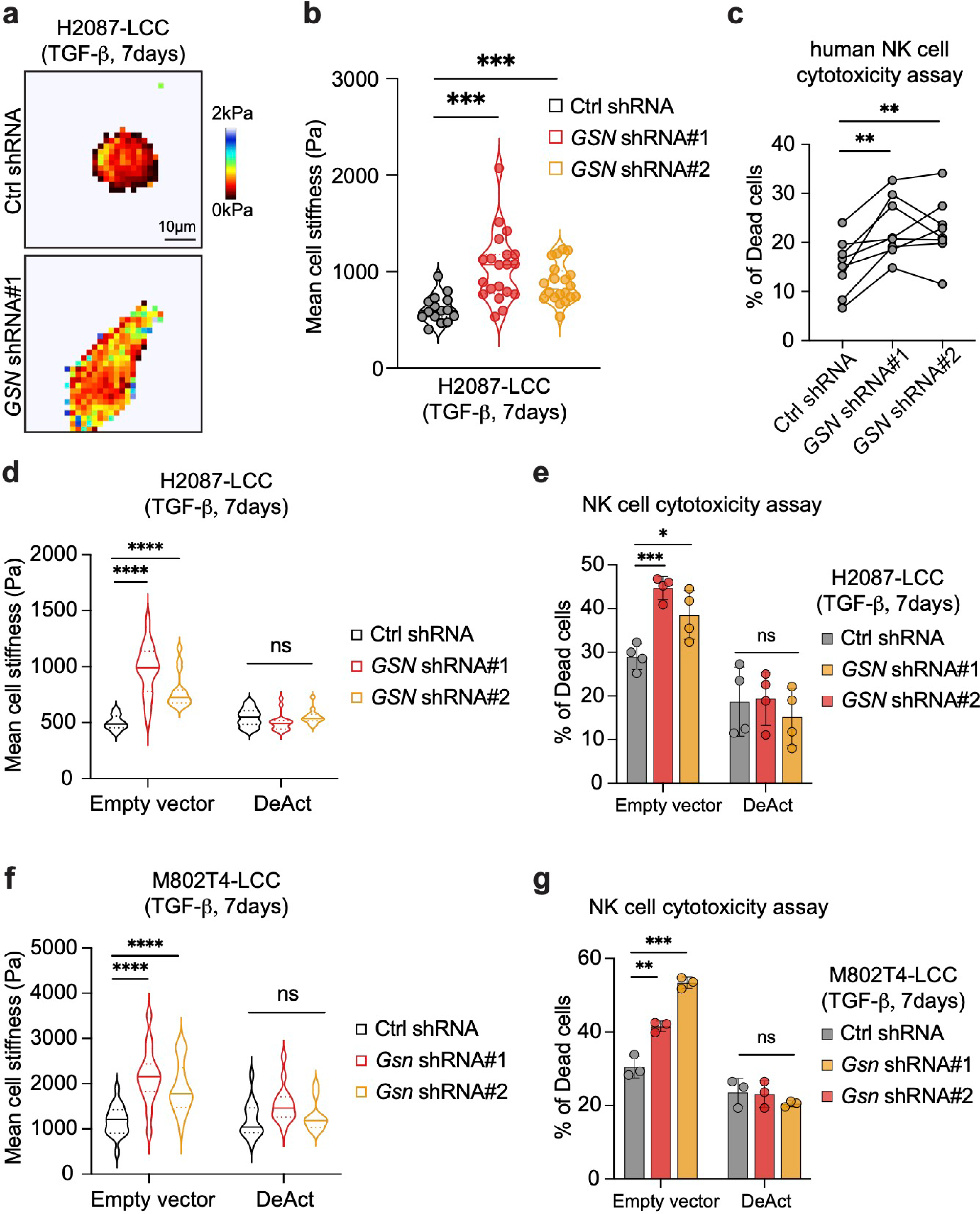
Gelsolin protects LUAD progenitors from mechanosurveillance. **a**, Force maps of representative H2087-LCC cells harboring control or *GSN-*targeting shRNAs and incubated with TGF-β for 7 days. **b**, Mean cell stiffness of H2087-LCC cells harboring the indicated shRNAs and incubated with TGF-β treatment for 7 days. n=18 per condition; two-sided unpaired *t*-test. **c**. cytotoxic effect of human NK cells on H2087-LCC cells harboring the indicated shRNAs and incubated with TGF-β for 7 days. Connecting lines indicate samples derived from the same donor (n=8). Paired *t*-test. **d**, Mean cell stiffness of H2087-LCC cells harboring the indicated shRNA and expressing control vector or DeAct. The cells were incubated with TGF-β treatment for 7 days. n=15 per condition, two-sided unpaired *t*-test. **e**, cytotoxic effect of mouse NK cells on H2087-LCC cells harboring the indicated shRNA and expressing control vector or DeAct. The cells were incubated with TGF-β for 7 days. Data are mean ± SEM, n=4; two-sided unpaired *t*-test. **f**, Mean cell stiffness of M802T4-LCC cells harboring the indicated shRNA and expressing control vector or DeAct. n=15 per condition, two-sided unpaired *t*-test. **g**, cytotoxic effect of mouse NK cells M802T4-LCC cells harboring the indicated shRNA and expressing control vector or DeAct. The cells were incubated with TGF-β for 7 days. Data are mean ± SEM, n=3; two-sided unpaired *t*-test.

To explore the role of gelsolin in mediating immune evasion of dormant metastatic cells in vivo, we intravenously inoculated H2087-LCC cells bearing control shRNA or shRNA against *GSN* into immunodeficient NSG mice. We observed no effect of gelsolin knockdown on lung colonization, indicating that gelsolin is not important for metastatic outgrowth in the absence of immune surveillance (Fig.7a). We then inoculated these cells into athymic mice and determined the number of disseminated cancer cells in the lungs. Gelsolin knockdown did not significantly alter the number of cancer cells disseminated to the lungs one week after injection, indicating that gelsolin is not essential for metastatic seeding in the lungs of a host that contains NK cells (Fig.7b). Of note, the knockdown of gelsolin decreased the number of H2087-LCC cells surviving in the lungs 7 weeks post injection (Fig.7b). Depleting NK cells in athymic mice by administrating anti-asialo-GM1 antibody 4 weeks after inoculation resulted in the formation of metastatic colonies by H2087-LCC cells harboring a control shRNA, implying that LCC cells withstand NK cell mediated killing in the dormant state and can subsequently grow after this immune pressure is relieved. By contrast, no metastases emerged after NK depletion in mice inoculated with gelsolin knockdown of H2087-LCC cells (Fig.7c), indicating that gelsolin is required for resistance to immune surveillance by NK cells during the dormant phase.

**Figure 7.**
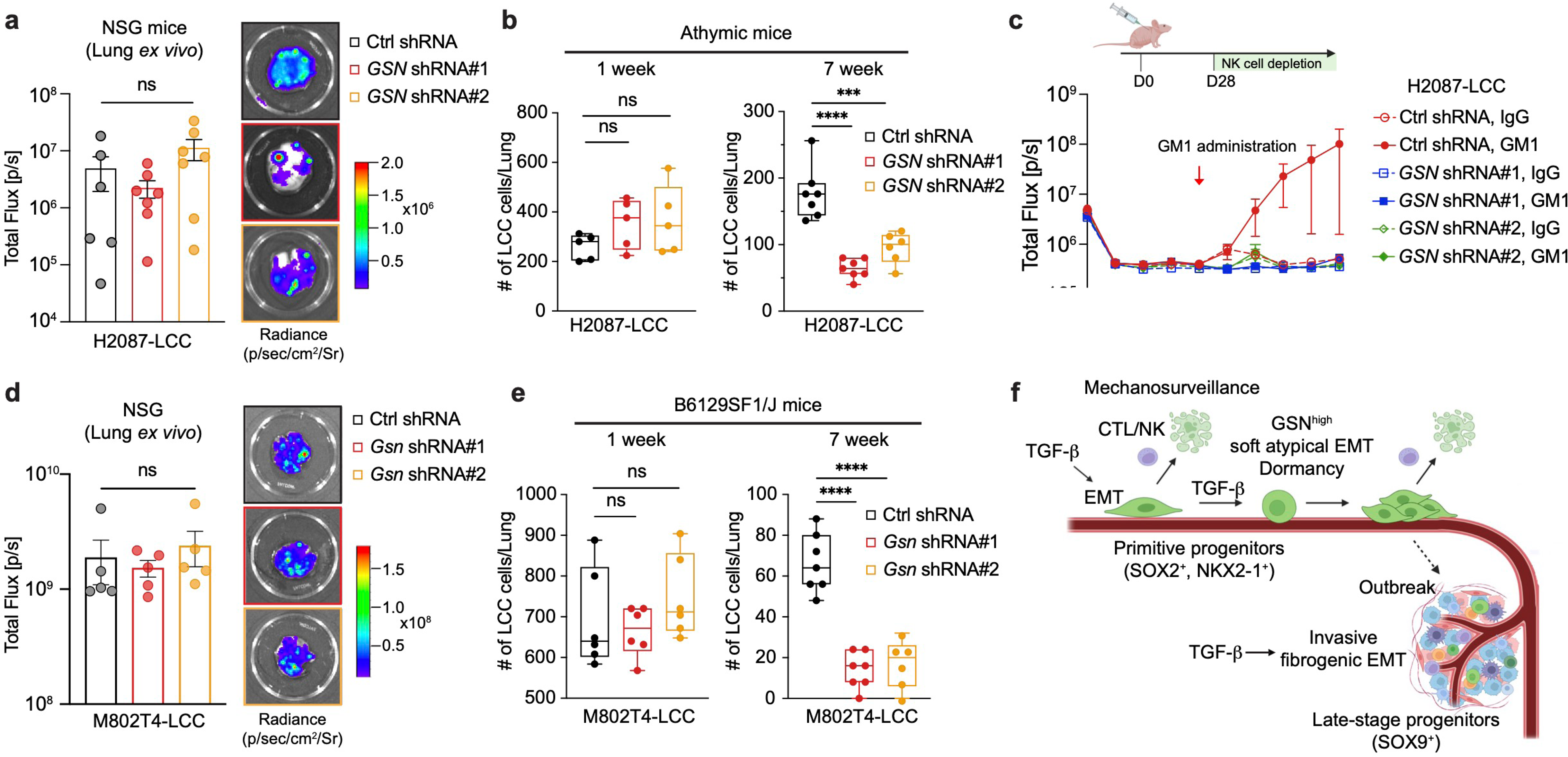
Gelsolin protects dormant metastasis from mechanosurveillance. **a**, Ex vivo BLI signal and representative images of lungs from NSG mice that were intravenously injected with H2087-LCC cell expressing the indicated shRNAs. Data are mean ± SEM; n=6 or 7 mice per group; two-sided Mann-Whitney U-test. **b**, Quantification of residual H2087-LCC cells in the lungs of athymic mice 1 week and 7 weeks after intravenous inoculation. n=5 or 7 mice per group; two-sided unpaired *t*-test. **c**. H2087-LCC cells expressing shRNA against GSN were intravenously inoculated into athymic mice. The mice were administered anti-asialo-GM1 antibody to deplete NK cells and allow the metastatic outgrowth of remaining disseminated cancer cells. IgG control or GM1 antibody administration started 4 weeks after cancer cell inoculation. Tumor burden was tracked by BLI analysis. Data are mean ± SEM, n=6 or 7 mice per group. **d**, Ex vivo BLI signal and representative images lungs from NSG mice that were intravenously injected with M802T4-LCC cell expressing the indicated shRNA. Data are mean ± SEM; n=5 mice per group; two-sided Mann-Whitney U-test. **e**, Quantification of residual M802T4-LCC cells in the lungs 1 week and 7 weeks after intravenous inoculation into B6129SF1/J mice. n=6 or 7 mice per group; two-sided unpaired *t*-test. **f**. Schematic summary of how TGF-β protects LUAD primitive progenitors (SOX2^+^ or NKX2-1^+^) cells from immune mechanosurveillance through gelsolin-modified EMT program (created with Biorender.com).

Similarly, knocking down gelsolin had no effect on lung colonization by M802T4-LCC in NSG mice (Fig.7d) or the number of these cells disseminated to the lungs one week after injection into immunocompetent mice, but strongly decreased the number of disseminated M802T4-LCC cells surviving in the lungs long-term (Fig.7e). Taken together, the evidence suggests that LUAD metastasis-initiating progenitor cells express gelsolin to prevent the accumulation of actin stress fibers during TGF-β-mediated dormancy, thereby diminishing cell stiffness and preventing effective mechanosurveillance by cytotoxic lymphocytes during long-term dormancy.

## Discussion

During the active colonization of new organs, cancer cells are thought to adopt a stiffer and migratory configuration that enables their invasion of perivascular niches, but which also sensitizes them to killing by CTLs and NK cells^33^. Focusing on the immediate stage after extravasation, our results show that LUAD primitive progenitors entering quiescence in response to TGF-β initially display the morphological, transcriptional, cytoskeletal, and migratory traits of a typical full EMT but subsequently transition into a soft, poorly migratory phenotype retaining a strong EMT transcriptional program but lacking actin stress fibers. This transition is mediated by a progressive increase in gelsolin expression which remodels the actin cytoskeleton. The resulting loss of stiffness in these cells allows them to evade immune mechanosurveillance and survive as metastatic progenitors during metastatic dormancy (Fig.7f).

Our results indicate that entering dormancy by metastatic cells is not a passive response to growth inhibitory cues in target organs. LUAD early progenitors appear to navigate this crucial step in disease progression by limiting the temporal scope of their mechanical vulnerability. It has been shown that softer cancer cells are more resistant to immune-mediated clearance both in vitro and in vivo^33,59,63^. The rapidity and prevalence of cell rounding during TGF-β-induced dormancy highlights the centrality of mechanosurveillance, and its evasion for the survival of dormant disseminated LUAD cells. It will be interesting to determine whether gelsolin-mediated cell softening or a functionally analogous mechanism contributes to dormancy in other cancer cell types. Of note in this regard, the gelsolin family contains additional isoforms (villin, advillin, capG) that have been linked to tumorigenesis and EMT^64-66^.

The atypical EMT induced by TGF-β in SOX2^+^ and NKX2-1^+^ LUAD cells contrasts with the EMT induced by TGF-β in cell populations derived from aggressive LUAD metastatic lesions, where SOX9^+^ late-stage progenitors are predominant. These populations, which are low in SOX2^+^ and NKX2-1^+^ cells, respond to TGF-β with a typical EMT coupled with expression of fibrogenic factors that support metastatic outgrowth^35,52^. Our findings suggest that as metastasis-initiating cells evolve, cells at different stages in a developmental continuum deploy distinct EMT programs in response to TGF-β to fulfill different pro-metastatic functions. In SOX2^+^ and NKX2-1^+^ LUAD primitive progenitors, TGF-β induces a dormancy-associated atypical EMT for evasion of immune surveillance, whereas in late-stage progenitors TGF-β induces a typical EMT coupled with microenvironment-modifying effectors for metastatic outgrowth^35,52^. In line with this idea, a previous study has identified multiple tumor subpopulations associated with different EMT and tumorigenicity levels in primary tumors^67^.

During each stage of the metastatic process, disseminated cancer cells are subjected to dramatic biomechanical adaptations to survive under extreme physical stress. Our findings support the concept that TGF-β-induced dormancy is a proactive process for the survival of metastatic progenitors. Besides downregulation of MHC molecules^19-22^, NK cell ligands^13^, and STING signaling^16^, our finding of gelsolin-mediated evasion of mechanosurveillance unexpectedly reveals a distinct strategy for metastatic progenitors to alleviate a liability associated with a full mesenchymal morphology, protecting these cells during the highly vulnerable period of solitary persistence^3,24^. These observations illustrate the complex dynamics of TGF-β in modeling EMT responses to ultimately promote metastasis. Importantly, the developmental stage of cancer cells is critical to enable TGF-β-induced immune evasion. We provide proof of principle that treatments that diminish this escape cause depletion of residual disease in our models, suggesting a basis for new approaches to adjuvant therapy.

## Acknowledgements

We thank the MSKCC Flow Cytometry Core Facility, Integrated Genomic Operation, and Y.H. Kim, W. Kang, Y. Romin and E. Chan of the MSKCC Molecular Cytology Core Facility for their technical assistance. We thank S. Malladi for the initial observation of the phenomenon investigated here, K. Srpan for human NK cells, and E.E. Er, J. Hu, and S. Gan for helpful discussions. This work was supported by National Institutes of Health grants R35-CA252978 (J.M.), P01-CA129243 (J.M.), R01-AI087644 (M.H.), R01-CA266068 (J.X.), P30-CA008748 (MSKCC), a grant from the Alan and Sandra Gerry Metastasis and Tumor Ecosystems Center at MSKCC (J.M.), T32 CA254875, a National Cancer Institute research training grant (Y.E.) and a postdoctoral fellowship from the Alan and Sandra Gerry Metastasis and Tumor Ecosystems Center (Z.W.).

## Author Contributions

Z.W. and J.M. conceived studies, designed experiments, interpreted results, and wrote the manuscript. Z.W. performed experiments and analyzed data. Y.E. and M.H. designed and performed CTL killing and activation assay and analyzed data. L.P. and G.L. provided technical assistance with experiments. Y.C. and J.X. designed and performed cell motility assay and analyzed data. I.G. performed IF staining of p27 expression in M802T4-LCC cells. M.H. provided assistance with manuscript preparation.

## Methods

### Animal studies

All animal experiments were performed in accordance with protocols approved by the Memorial Sloan Kettering Cancer Center Institutional Animal Care and Use Committee (IACUC). Athymic nude mice were obtained from Envigo (strain #: 069) or Charles River Laboratories (strain #: 490). NSG (NOD.Cg-Prkdc^scid^IL2rg^tm1Wjl^/SzJ, strain #005557), B6129SF1/J (strain #101043) and B6(Cg)-*Tyr^c-2J^*/J (B6-albino, strain #000058) mouse strains were obtained from the Jackson Laboratory. Female mice 6 to 8 weeks of age were used for in vivo studies. 2 to 6 months-old male and female OT1 αβTCR transgenic mice (Jackson Laboratories, Strain #:003831) were used to generate OT1 CTLs for in vitro assays. All animals were housed under specific pathogen-free conditions.

For brain metastasis assays, 1x10^5^ cells were resuspended in 100 µl of phosphate-buffered saline solution (PBS) and intracardially injected into the right ventricle with a 26G tuberculin syringe. Lung colonization assays were performed by injecting 1x10^5^ or 5x10^4^ cells in the lateral tail vein of mice with a 28G insulin syringe. Metastatic burden was monitored by bioluminescence imaging (BLI) using retro-orbital injection of D-luciferin (150 mg/kg) and an IVIS Spectrum Xenogen instrument (PerkinElmer). Data were analyzed using Living Image software v4.5 (PerkinElmer). For ex vivo imaging of metastases-bearing organs at the experimental end point, animals were anesthetized with 100 mg/kg of ketamine and 10 mg/kg of xylazine and retro-orbitally injected with D-luciferin prior to organ isolation. Isolated organs were analyzed using an IVIS Spectrum Xenogen instrument and Living Image software.

For experiments with cells engineered with a TGF-β reporter, mice were maintained on regular diet and switched to a diet of doxycycline food pellets (2,500 mg kg−1, Envigo) two weeks prior to harvesting organs for analysis.

For NK cell depletion experiments in athymic nude mice, 33 μg of anti-asialo-GM1 antibody (Wako Chemical, Cat# 986-10001) was injected intraperitoneally per mouse once every 5 days. For NK, CD4 or CD8 T cell depletion assays in C57BL/6J and B6(Cg)-*Tyr^c-2J^*/J mice, 200 μg *InVivo*Mab anti-mouse NK1.1 antibody (clone PK136, BioXCell, Cat# BE0036), CD4 antibody (clone GK1.5, BioXCell, Cat# BE0003-1), CD8α antibody (clone 53-6.7, BioXCell, Cat# BE0004-1), or IgG2a control (clone 2A3, BioXCell, Cat# BE0089) was injected intraperitoneally in each mouse once weekly. Mice were randomly assigned to control or treatment groups. To validate the depletion of each immune population, peripheral blood was collected and stained with violetFluor 450 Anti-Mouse CD45 (30-F11) (Tonbo Biosciences, Cat# 75-0451-U100), FITC Anti-Mouse CD4 (GK1.5) (Tonbo Biosciences, Cat# 35-0041-U100), PE-Cy7 Anti-Mouse CD8a (53-6.7) (Tonbo Biosciences, Cat#60-0081-U100), APC Anti-Mouse NK1.1 (CD161) (PK136) (Tonbo Biosciences, Cat# 20-5941-U100), followed by flow cytometric analysis on an LSRFortessa (BD Biosciences) instrument.

### Cell culture

H2087-LCC cells and derivatives from spontaneous metastatic outbreaks (SO) of H2087-LCC cells were cultured in RPMI 1640 media supplemented with 10% fetal bovine serum (FBS), 2 mM glutamine, 100 IU/mL penicillin/streptomycin, 1 µg/mL amphotericin B, 0.5 mM sodium pyruvate, 10mM HEPES, 50 nM hydrocortisone, 25 nM sodium selenite, 20 μg/mL insulin, 10μg/mL transferrin (SITE), 0.5% bovine serum albumin (BSA), and 1 ng/mL recombinant human epidermal growth factor. Mouse lung cancer cell line 802T4 and 393T3 was a gift from T. Jacks (Kock Institute MIT) ^36^. M802T4-LCC, spontaneous outbreak derivatives (M802T4-SO), 393T3 and A549 cells were cultured in RPMI 1640 media supplemented with 10% FBS, 2 mM glutamine, 100 IU/mL penicillin/streptomycin, and 1 µg/mL amphotericin B. 293T cells were cultured in DMEM supplemented with 10% FBS and 2 mM glutamine. All cell lines tested negative for mycoplasma contamination.

For isolation of M802T4-LCC cells, 802T4 cells (1x10^5^) expressing a lentiviral vector encoding firefly luciferase and green fluorescent (GFP) construct in combination with puromycin antibiotic resistance marker in a volume of 100 µL was intravenously injected into the lateral tail vein of B6129SF1/J mice. Cancer cell colony growth was tracked by BLI. Lungs from BLI negative mice were resected under sterile conditions and dissociated using Lung Dissociation Kit (Miltenyl Biotec, Cat# 130-095-927) following the manufacturer’s instructions. Cells were then resuspended in culture conditions and allowed to attach on a 15-cm dish. Cancer cells from these BLI-negative lungs were selected with 2.5 μg/ml of puromycin (Gibco). For establishing cell lines from spontaneous outbreaks, animals with BLI detectable metastases were killed and organs were imaged ex vivo to confirm the presence of macrometastatic lesions. Organs were resected under sterile conditions and mechanically dissociated using a gentleMACS dissociator (Miltenyi Biotec) and placed in culture medium containing a 1:1 mixture of DMEM/Ham’s F12 medium supplemented with 0.125% of collagenase III and 0.1% of hyaluronidase. Minced samples were incubated at 37 °C for 1 h, with gentle rocking to produce single-cell suspensions. Cells were then resuspended in their respective culture conditions and allowed to grow to confluence on a 15-cm dish. Cancer cells were selected by adding 5 μg/ml of blasticidin (H2087 cells) or 2.5 μg/ml of puromycin (M802T4 cells) to the media.

For analysis of TGF-β responses, cells were cultured in media containing 2% FBS and incubated with SB-505124 (2.5μM, Millipore Sigma, Cat# S4696-5MG) or 100 pM TGF-β1 (R&D Systems) for the indicated period of time. SB-505124 inhibits the kinase activity of the TGF-β subfamily type 1 receptors TGFBR1 (also known as ALK5), ACVR1B (ALK4), and ACVR1C (ALK7).^47^ Bright-field images were acquired with EVOS M5000 microscope (Thermo Fisher Scientific).

For analysis of Wnt-3a response, cells were cultured in media containing 2% FBS and treated with recombinant human Wnt-3a protein (R&D systems, Cat# 5036-WN-010) at a final concentration of 200ng/mL for 2h.

### Gene knockout, knockdown and overexpression constructs

CRISPR-mediated knockouts were generated by cloning sgRNAs into the Guide-it CRISPR/Cas9 vector (Red, TaKaRa, Cat# 632602), transfecting the construct into cells, and isolating and expanding the cells with knockouts from single-cell colonies. Sequences of sgRNA oligos are listed below:

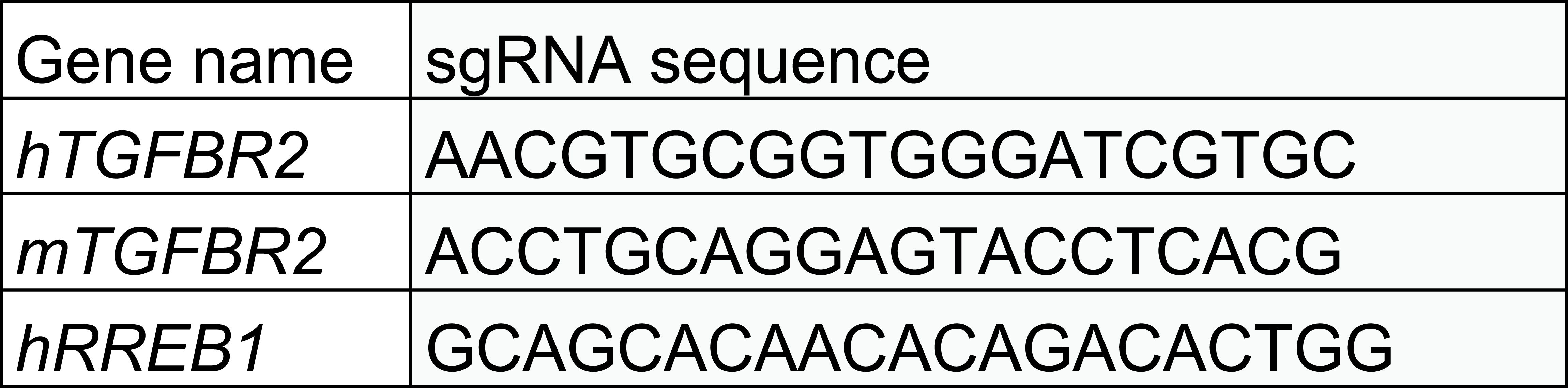

Lentivirus suspensions were produced by transfection of lentiviral vector with second generation packaging constructs psPAX2 and pMD2.G (Didier Trono, Addgene plasmids 12260 &12259) into 70% confluent 293T cells using Lipofectamine 2000. Viral particles were collected, filtered through 0.45 μm sterile filters, and incubated with the cells of interest for 12 h with 8 μg/mL polybrene. Cells were recovered in growth medium overnight before addition of selection media including 500 μg/mL hygromycin (Life Technologies, 10687-010), 10 μg/mL puromycin (Sigma).

Stable knockdowns of *GSN* in H2087-LCC cells were generated using two independent shRNA purchased from Millipore Sigma (Mission Predesigned shRNA clone, shRNA#1: TRC Clone ID# TRCN0000029725; shRNA#2: TRC Clone ID# TRCN0000029728). Non-mammalian shRNA was used as control (Millipore Sigma, Cat# SHC002). Stable knockdowns of *Gsn* in M802T4-LCC cells were generated using two independent shRNA purchased from Horizon Discovery (SMARTvector Lentiviral shRNA, shRNA#1: Clone ID# V3SVMM08_10907046; shRNA#2: Clone ID# V3SVMM08_11757555). Non-targeting shRNA was used as control (Horizon Discovery, Cat#: VSC11715)

For DeAct-SpVB expression, the DHRFdd-SpvB (Salmonella SpvB 375-591) sequence was amplified by PCR from pTetON-DHFRdd-SpvB; CMV-mCherry (Addgene, plasmid # 89463). PCR amplified products were ligated to XmalI/MluI-digested pLVX-TRE3G-mCherry (Takara, Cat #: 631360), a tetracycline-inducible lentiviral expression vector. For all the experiments, expression of DeAct was induced 24h before analysis.

### Immunofluorescence, live imaging and EdU cell proliferation assay

For immunofluorescence (IF) staining of tissue samples, organs were fixed in 4% paraformaldehyde overnight at 4°C and then washed twice with PBS. Organs were cryo-protected by immersion in 30% sucrose, then mounted using OCT (Sakura, Cat# 4583) on a sliding microtome with a platform freezing unit (Thermo Fisher Scientific, Cat# Microm KS-34 and Microm HM-450). 80 μm sections were cut and stored in anti-freezing solution (30% v/v ethylene glycol, 30% glycerol v/v in PBS) at -20°C. Floating sections representative of the entire organ were permeabilized by washing in 0.25% Triton X-100 in PBS with (PBS-Tr) three times, followed by incubation for 1 h in blocking buffer containing 10% normal goat serum (Life Technologies, Cat# 50062Z), 2% BSA (Fisher Scientific, Cat# BP9706100), and 0.25% Triton X-100. Sections were then incubated with primary antibodies diluted in blocking buffer, overnight at 4°C. After washing with PBS-Tr six times, sections were incubated in fluorophore-conjugated secondary antibodies for 2 h. Sections were washed with PBS-Tr and then PBS three times each, followed by staining with 4’,6-diamidino-2-phenylindole (DAPI) nuclear dye (Thermo Fisher Scientific, Cat# D3571) for 5 min, and three additional washes with PBS. Sections were transferred onto slides and mounted using ProLong diamond antifade mountant (Life Technology, Cat# P36970).

For immunofluorescence staining of cells in culture, cell plated in Nunc Lab-Tek II chamber slides (Thermo Fisher, Cat#154453) were fixed for 15 min at room temperature in 4% paraformaldehyde and then washed three times with PBS. Samples were incubated for 1 h in blocking buffer containing 10% normal goat serum (Life Technologies, Cat# 50062Z), 2% BSA (Fisher Scientific, Cat# BP9706100), and 0.25% Triton X-100. Cells were then incubated in primary antibodies, diluted in blocking buffer, overnight at 4°C or for 2 h at room temperature. After washing with PBS three times, cells were incubated in fluorophore-conjugated secondary antibodies for 1 h at room temperature. Cells were washed with PBS three times, followed by staining with DAPI (Thermo Fisher Scientific, Cat# D3571) for 5 min. Slides were mounted using ProLong diamond antifade mountant (Life Technology, Cat# P36970).

Immunofluorescent images were acquired with a Zeiss Axio Imager Z1 microscope or an SP5 confocal microscope (Leica Microsystems). Fluorescent intensity was analyzed with ImageJ software. Antibodies used for immunofluorescence are listed below:

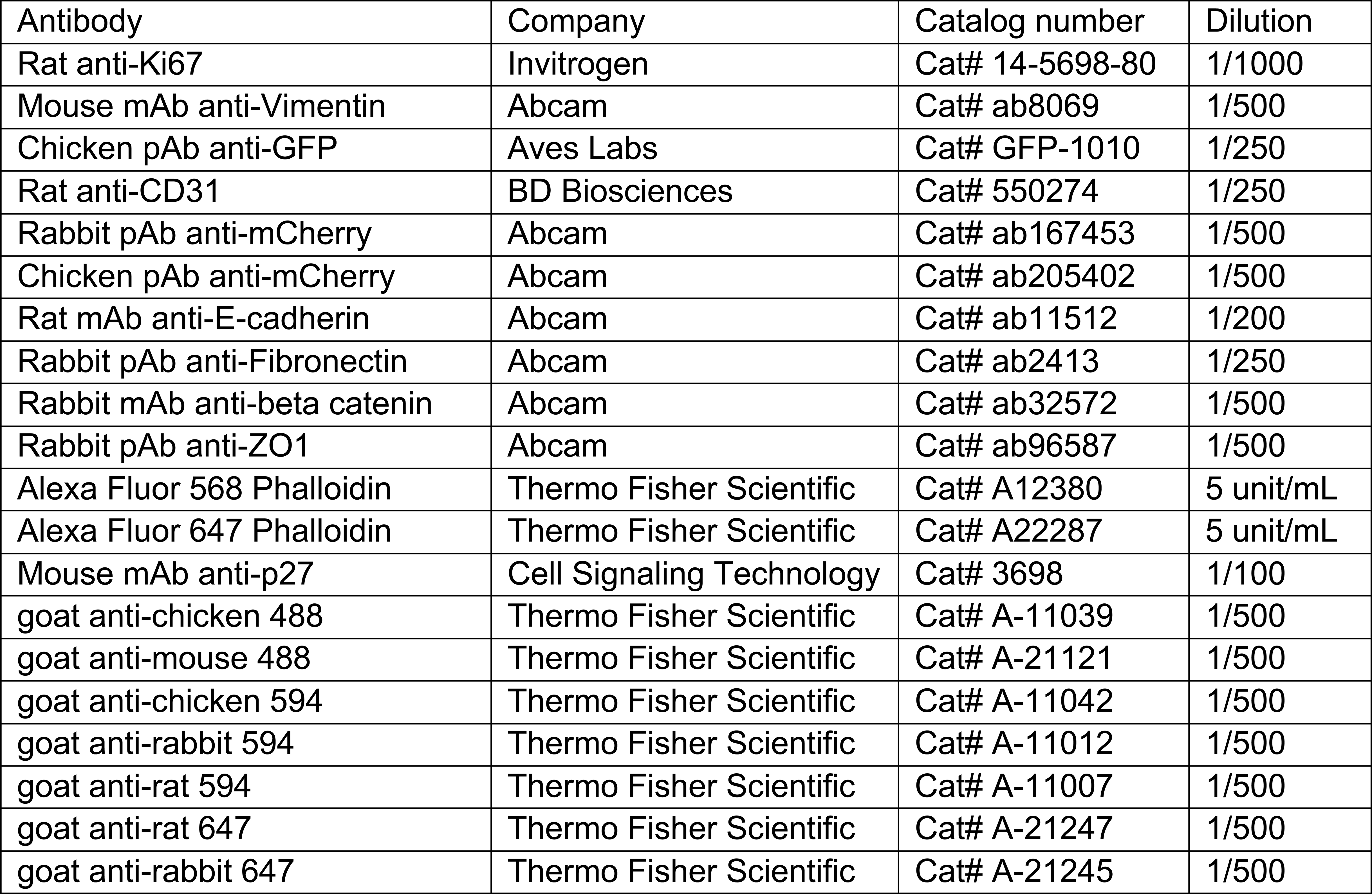

For time-lapse live imaging, 2x10^4^ cancer cells were plated in 6-well plates and incubated with TGF-β for 5 days. Plates were imaged at 6 min intervals using a Zeiss Zen epifluorescence microscope at 10x magnification for 48 h. For live imaging of NK cell-mediated killing, cancer cells were incubated with TGF-β for 5 days and labeled with CellTracker Green CMFDA Dye (Thermo Fisher Scientific, Cat# C7025). After washing, cells were mixed with mouse NK cells at a 1:4 carcinoma to NK cell ratio, in the presence of 1.5 µM propidium iodide. Plates were imaged at 6 min intervals using a Zeiss Zen epifluorescence microscope at 10x magnification for 5 h.

For EdU cell proliferation assay, cells were plated in Nunc Lab-Tek II chamber slides (Thermo Fisher, Cat#154453), followed by indicated treatments. EdU incorporation assay was performed with Click-iT™ Plus EdU Cell Proliferation Kit for Imaging, Alexa Fluor™ 594 dye (Thermo Fisher Scientific, Cat# C10639). Cells were incubated with 10 μM EdU for 1h, following the manufacturer’s instructions. 15-18 images were collected for each condition using a 40× objective on a Zeiss Axio Imager Z1 microscope. Fluorescence intensity was analyzed with ImageJ software (v2.14.0).

### RNA sequencing and data analysis

Total RNA purified from cells was quantified by Ribogreen and quality assessed by Agilent BioAnalyzer. 500 ng of RNA with integrity number (RIN) > 9.5 from each sample was used for library construction with TruSeq RNA Sample Prep Kit v2 (Illumina) according to manufacturer’s instructions. Multiplexed sequencing libraries were run on a Hiseq2500 platform and more than 30-40 million raw paired-end reads were generated for each sample. For data analysis, reads pairs in FASTQ format (50bp/50bp) were assessed for quality using FastQC v0.11.5 and mapped to human genome hg19 with STAR2.5.2b using standard settings for paired reads. Uniquely mapped reads were assigned to annotated genes with HTSeq v0.6.1p1 with default settings. Read counts were normalized by library size, and differential gene expression analysis based on a negative binomial distribution was performed using DESeq2 v3.4. Gene set enrichment analysis (GSEA) was performed using previously curated gene sets.

### qRT-PCR analysis

RNA was extracted from cells using the RNeasy Mini Kit (Qiagen, Cat# 74106). cDNA was generated using the Transcriptor First Strand cDNA Synthesis Kit (Roche, 04379012001). Relative gene expression was determined using Taqman assays (Life Technologies) or SYBR green assays (Life Technologies). Quantitative PCR was performed on the ViiA 7 Real-Time PCR System (Life Technologies). Housekeeping gene GAPDH was used as the internal control for calculating relative gene expression.

Taqman assays used for human genes were *GAPDH* (Hs99999905_m1), *GSN* (Hs00609272_m1), *HAS2* (Hs00193435_m1), *IL11* (Hs01055413_g1), *SERPINE1* (Hs00167155_m1), *SNAI1* (Hs00195591_m1), *DKK1* (Hs00183740_m1) and *Axin2* (Hs00610344_m1). Taqman assays used for mouse genes were: *Gapdh* (Mm99999915_g1), *Snai1* (Mm00441533_g1) and *Gsn* (Mm00456679_m1). Primers for SYBR green assays were:

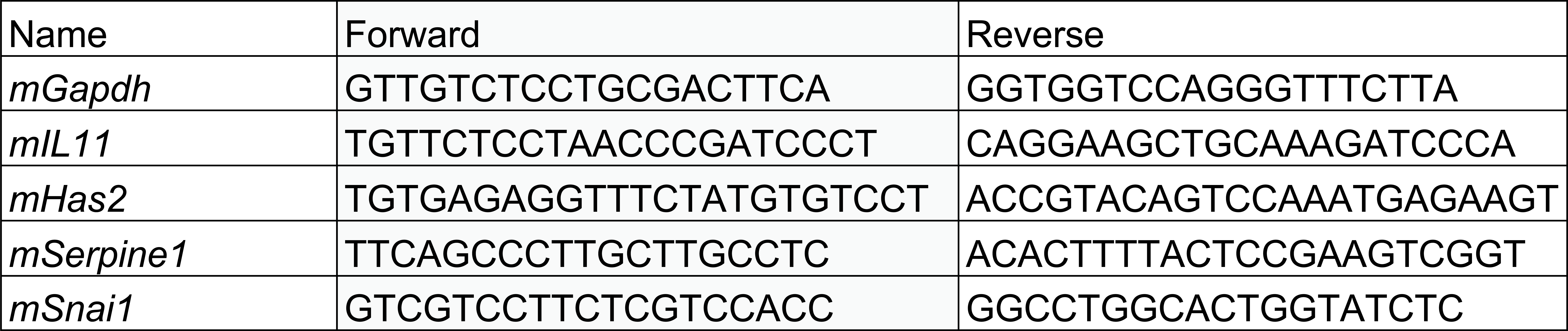

### Chromatin immunoprecipitation (ChIP) PCR analysis

For ChIP, cells were crosslinked with 1% formaldehyde (Millipore Sigma, Cat# F8775) for 10 min and quenched with 0.125 M glycine for 5 min at room temperature. ChIP was performed using an assay kit (Millipore Sigma, Cat# 17-295) according to the manufacturer’s instructions. Antibodies against H3K27Ac (Active Motif, Cat# 39133, 5 µg per sample), H3K27me3 (Millipore Sigma, 07-449, 10 µg per sample), H3K4me1(Abcam, ab8895, 5 µg per sample) were used. Immunoprecipitated DNA was purified using the QIAquick PCR Purification Kit (Qiagen), followed by qPCR analysis. The amplification product was calculated as the percentage of the input, then normalized to the control experiment for each condition. ChIP-PCR primers are listed below:

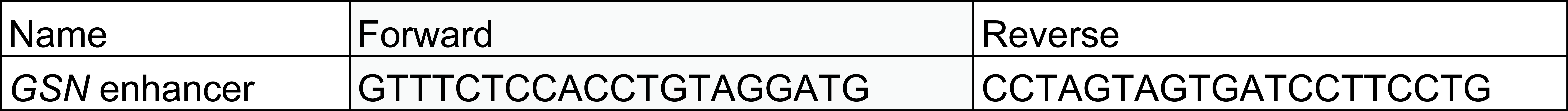

### Western immunoblotting and ELISA

Cells were lysed using RIPA cell lysis buffer (Cell Signaling Technology, Cat# 9806S) supplemented with a protease inhibitor cocktail (Roche, cOmplete, mini, EDTA-free protease inhibitor tablets, Cat# 11836170001) and a phosphatase inhibitor cocktail (Thermo Fisher Scientific, Halt Phosphatase Inhibitor Cocktail, Cat# 78427, 1:100). Protein concentration was measured using the Pierce BCA Protein Assay Kit (Thermo Fisher Scientific, Cat# 23227). Protein was then mixed with NuPage LDS sample buffer (Thermo Fisher Scientific, Cat# NP0007) supplemented with NuPAGE Sample Reducing Agent (Thermo Fisher Scientific, Cat# NP0009), heated at 95°C for 10 min, then loaded in NuPAGE Novex 4%-12% Bis-Tris gels (Thermo Fisher Scientific, Cat# NP0336BOX) using 1x MOPS SDS running buffer (Thermo Fisher Scientific, Cat# NP0001) and transferred to nitrocellulose membranes. Membranes were blocked with Odyssey blocking buffer (LI-COR Biosciences, Cat# 927-60001) and incubated overnight at 4°C with primary antibodies diluted in blocking buffer. After washing with PBS containing 0.1% Tween (PBST) three times, membranes were incubated with IRDye 680RD goat anti-mouse (LI-COR Biosciences, Cat# 926-68070, 1:10000), IRDye 680RD goat anti-rat (LI-COR Biosciences, Cat# 926-68076, 1:10000) or IRDye 800CW goat anti-rabbit (LI-COR Biosciences, Cat# 926-32211, 1:10000) secondary antibodies. Signal was detected using an Odyssey CLx imager (LI-COR Biosciences). Antibodies used for immunoblotting are listed below:

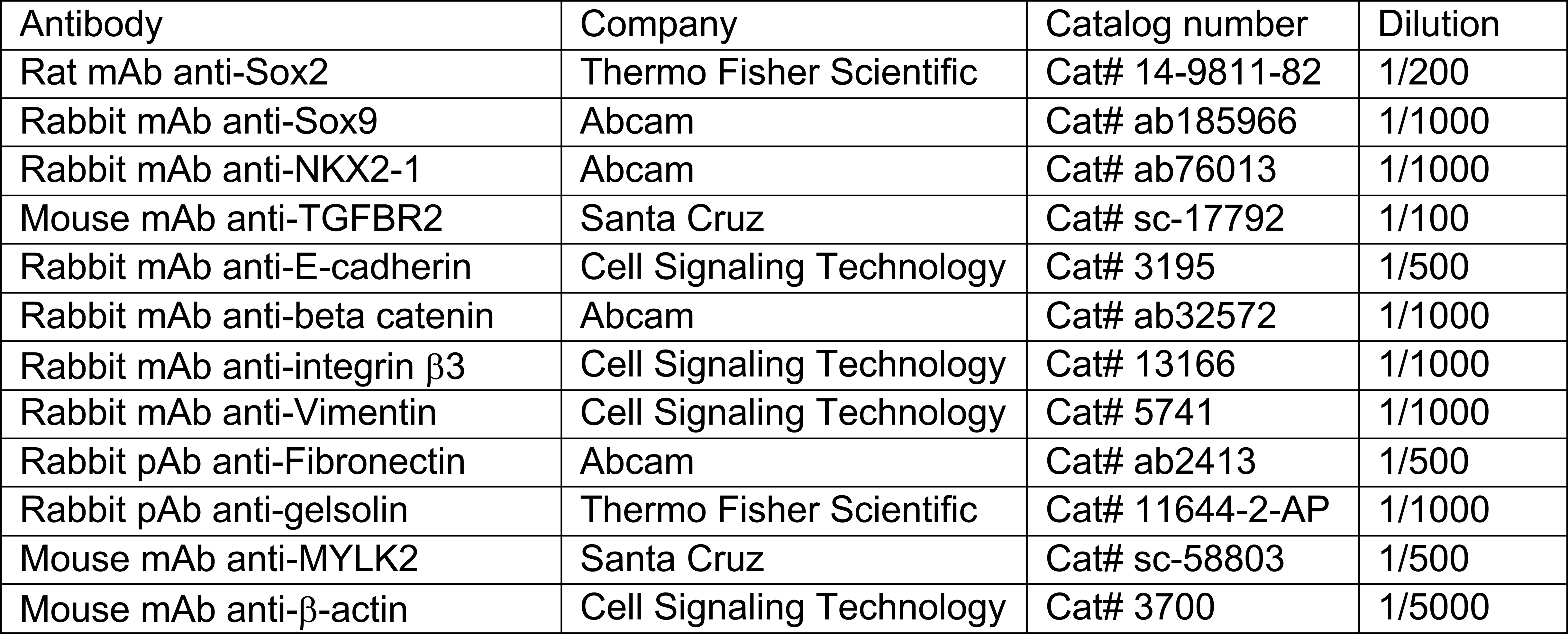

For determination of DKK1 protein level, 2×10^4^ cells were plated in 6-well plate, followed by indicated treatment. Fresh medium with SB-505124 or TGF-β was changed 24h before supernatant collection. DKK1 levels were measured using Human Dkk-1 Quantikine ELISA Kit (R&D systems, Cat# DKK100B), following the manufacturer’s instructions. Cell numbers were determined at the time of supernatant collection for normalization.

### Atomic Force Microscopy (AFM)

Cells were seeded on glass-bottom Petri Dishes (FluoroDish FD35) and kept in cultured medium during the acquisition of stiffness maps. AFM images were captured with an Nanowizard V (JPK-Bruker) in QITMadvanced Mode (stiffness mapping) at 37°C. For cell stiffness mapping, 1 µm diameter spherical AFM probe (silicon nitride cantilever, nominal spring constant k = 0.2 N/m, SAA-SPH-1UM, Bruker) was used. The spring constant of each AFM probe was measured by the thermal noise method. 15-20 cells from each experimental group were measured in each session. Bright field images of each cell were collected during AFM measurements in an inverted optical objective integrated with the AFM (Zeiss AxioObserver Z1). For the stiffness mapping, 1 nN setpoint was used (60 µm x 60 µm or 120 µm x 120 µm image size with 32 x 32 pixel resolution) to ensure 2-3 µm sample indentation. The data were processed with JPK Data Processing software using the Hertz model with 0.5 Poisson ratio as a fit parameter. Stiffness histograms were obtained by identifying the stiffness values belonging to each cell (and not the substrate values, shown in black on force maps) through a mask, and plotting the results from each cell sample as a single population. All measurements made <2µm above the substrate were excluded. Stiffness distribution histograms were obtained using the histogram analysis tool in Excel (Microsoft) after normalizing for the total number of data points.

### Lymphocyte cytotoxicity, degranulation, and cytokine production assays

To isolate murine NK cells, splenocyte suspensions were prepared by mechanical dissociation. NK cells were purified by magnetic depletion of non-NK cells using NK Isolation Kit II (Miltenyi Biotec, Cat# 130-096-892) and separation with magnetic columns (Miltenyi Biotec, Cat# 130-042-401). NK cells were cultured in NK cell medium (RPMI-1640 medium supplemented with 10% FBS, β2-mercaptoethanol, non-essential amino acids, 10 mM HEPES, 0.5 mM sodium pyruvate, 2 mM L-glutamine, and 10 IU/ml of penicillin/streptomycin) containing 1,000 U/ml of murine recombinant IL-2 (R&D Systems, Cat# 402-ML-020).

To isolate human NK cells, peripheral blood was collected from healthy donors using protocols approved by the Memorial Sloan Kettering Cancer Center Institutional Review Board (nos. 06-107 and 95-054). The samples were processed under Biospecimen Research Protocol Institutional Review Board no. 16-1564. Donors provided informed written consent. Peripheral blood mononuclear cells were isolated by Ficoll gradient purification. NK cells were purified using NK cell isolation kit (Miltenyi Biotec, Cat# 130-092-657) and expanded with C9 feeder cells at a 1:1 ratio in R10 media supplemented with 200 IU/ml human recombinant IL-2 (PeproTech, Cat# 200-02).

For measuring NK killing of cancer cells, target cancer cells plated in 3 or 4 replicates in 96-well plates were labeled with CellTracker Green CMFDA Dye (Thermo Fisher Scientific, Cat# C7025), and incubated with NK cells at a 1:4 carcinoma to NK cell ratio, for 4 h at 37 °C. Cell mixtures were stained with 7-AAD Viability Staining Solution (Thermo Fisher Scientific, Cat# 00-6993-50) to assess cancer cell cytolysis by flow cytometry.

For CTL cytotoxicity assays, cancer cells plated in triplicates in 96-well plates were loaded with varying concentrations of OVA peptide (1 nM, 0.3 nM, 0.01 nM, 0.03 nM, 0 nM) for 2 h and washed three times in culture medium. To assess killing, OT1 CTLs were added at a 4:1 effector to target ratio and incubated for 5 h at 37°C in culture medium. Cells were then labeled with allophycocyanin (APC) conjugated anti-CD8a antibody (Tonbo Biosciences, Cat# 20-0081), and specific lysis of target cells (GFP+; CD8-cells) was determined based on incorporation of either propidium iodide (Thermo Fisher Scientific, Cat# P3566) or DAPI (Invitrogen, D1306) using flow cytometry.

To assess lytic granule secretion, a 2:1 CTL:cancer cell ratio was used. Cells were incubated for 90 min at 37°C in the presence of eFluor660-conjugated anti-Lamp1 antibody (1 μg/ml, Clone 1D4B, eBiosciences). Cells were then labeled with anti-CD8a antibody, and the percentage of CTLs (CD8+ cells) with positive Lamp1 staining was quantified by flow cytometry. To assess cytokine production, 4:1 CTL:cancer cell admixtures were incubated for 4 h at 37°C in the presence of BD GolgiPlug protein transport inhibitor (BD Biosciences). Cells were then labeled with anti-CD8a antibody, fixed, and permeabilized using the BD Cytofix/Cytoperm kit. After labeling with FITC-conjugated anti-TNF (BioLegend 506304) and PE/Cy7-conjugated anti-IFNγ (BioLegend, 505826) antibodies, the percentage of CTLs (CD8+) expressing TNF and IFNγ was determined by flow cytometry. All assays were performed in triplicate.

### Cell motility assay

Time-lapse microscopy was performed on an inverted microscope (Zeiss AxioObserver Z1) using a 10x/0.45 NA objective. 2000 H2087-LCC cells were seeded in 6-well plate and treated with TGF-β for 3 days or 7 days prior to imaging. The temperature was set to 37°C in the incubation chamber (incubator XLmulti S1). Cells at six different positions were imaged every 30 min for 24 h. Both phase contrast images with 100 ms exposure time and fluorescent images (GFP, Ex: 488 nm, Em: 509 nm) with 20 ms exposure time were acquired. Excitation was performed using a LED lamp (Colibri 7). The images were recorded using a digital camera (Hamamatsu, c4742-98) at 0.645 μm/pixel resolution using 1345 pixel × 1025 pixel image sizes at 12 bits depth. The software for image acquisition was Zen 2.6.

Image preprocessing (contrast adjusting, background subtraction, segmentation) was performed using Fiji^68^. Centroids of cells were identified by our customized MATLAB script, and tracks were generated using a MATLAB implementation of IDL tracking methods developed by J. Crocker, D. Grier, and E. Weeks (https://site.physics.georgetown.edu/matlab/code.html). All tracks were reviewed by cross-comparison with centroid movies and removed manually when they were invalid. Cell types were categorized at the start into more rounded (type 1) and less rounded (type 0) types according to a threshold value of 0.6 in roundness.

For each cell track, mean squared displacement (MSD) was calculated over a range of lag times as previously described^69^. The log-log plot of MSD versus lag time provides information about both the diffusion coefficient (intercept) and persistence (slope, α) of cells. For a cell moving randomly, α=1; for a cell with applied force, α>1; and for a cell that is constrained, α<1. Motility (α) was quantified as the slope using the *fitlme* function in MATLAB with the model, ‘log(MSD) ∼ 1+ cellType*log(τ)’^70^. The model fits the 1/4 of log(MSD) points obtained from each trajectory as MSDs present large statistical fluctuations when τ is large^69^.

### Statistics

Statistical analysis details are described each figure legend. Mann-Whitney U-test, t-test, Chi-square test and log-rank test were performed in Graphpad Prism. The number of samples (n) is indicated in each figure panel or figure legend.

## Resource availability

## Lead Contact

Further information and requests for resources and reagents should be directed to and will be fulfilled by the lead contact Joan Massagué.

## Materials availability

All unique reagents generated in this study, including plasmids and cancer cell lines, are from the lead contact with a completed Materials Transfer Agreement.

## Data and code availability

Raw sequencing reads and processed files for RNA-seq have been deposited in the Gene Expression Omnibus database (GEO) under the SuperSeries accession number GEO: GSE269762 and are publicly available as of the date of publication. All software programs used for analyses are publicly available and described in the method section.

## Extended Data Figure Legends

**Extended Data Fig.1.**
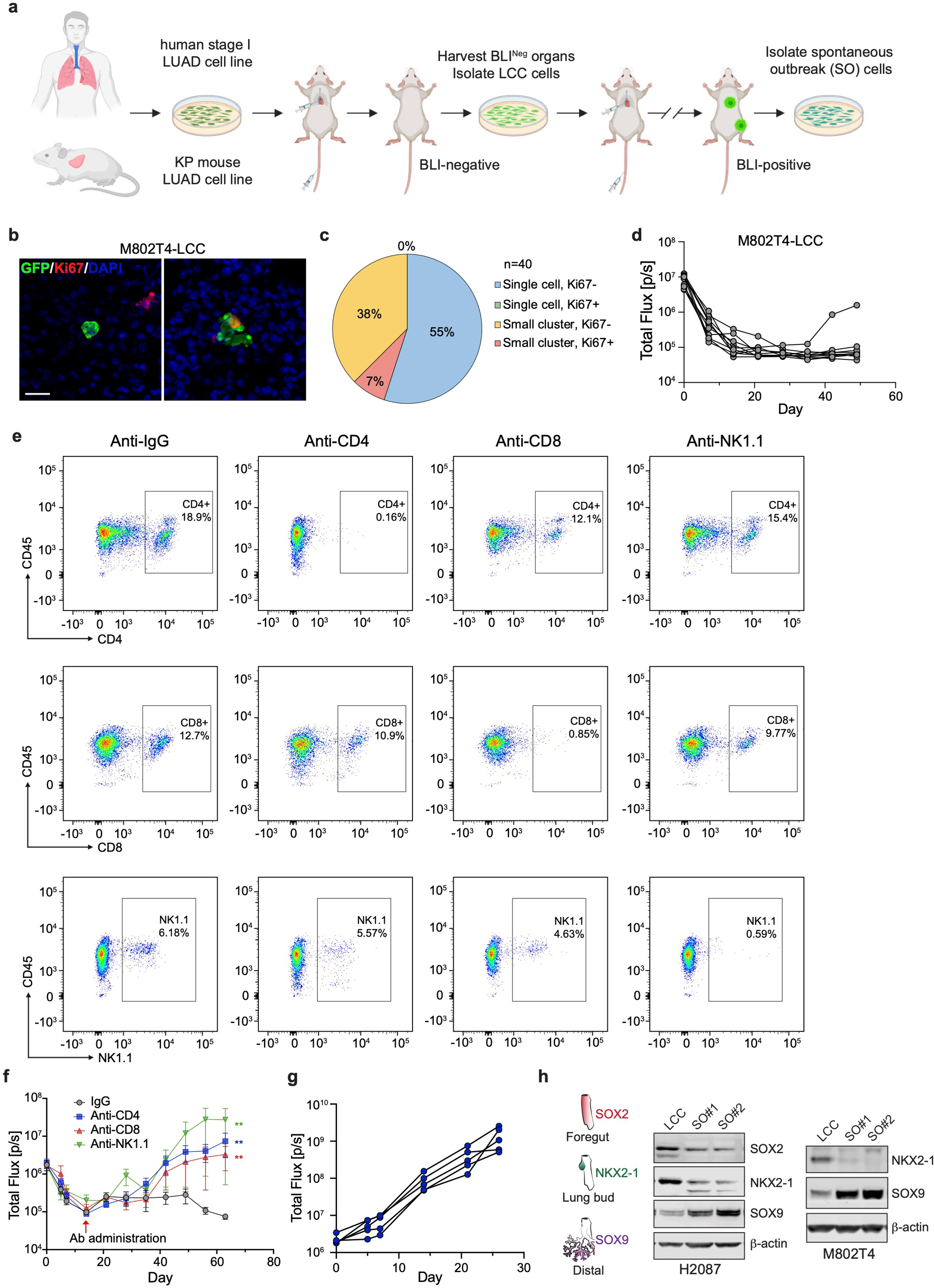
Generation and features of dormant LUAD metastasis models. **a**, Schematic protocol for the isolation of human or mouse LUAD cancer cells that are competent to enter dormancy and establish latent metastasis (created with Biorender.com). **b**, Representative images of a single cell or small cluster of M802T4-LCC cells in the lungs of B6129SF1/J mice 7 weeks after intravenous injection. The cells expressed green fluorescent protein (GFP), and firefly luciferase for bioluminescence imaging (BLI). Ki67 in red and DAPI in blue. Scale bar: 20 μm. **c**, Quantification of the proliferation status of M802T4-LCC cells and clusters (fewer than 20 cells) in the lungs 7 weeks after intravenous injection. n=40. **d**, BLI tracking the long-term persistence and metastatic outbreaks in the lungs of M802T4-LCC cells after intravenous inoculation into B6129SF1/J mice. 1 out of 10 mice developed overt metastases. **e**, Representative flow cytometry analysis of immune population in peripheral blood of B6129SF1/J mice injected with indicated antibody. **f**, BLI tracking of M802T4-LCC cells intravenously inoculated into B6129SF1/J mice followed by antibody-mediated depletion of the indicated immune cells. Data are mean ± SEM; n=6 or 7 mice per group; two-sided Mann-Whitney U-test. **g**, BLI tracking of M802T4-LCC tumor growth after intravenous injection into NSG mice. n=5 mice. **h**, Western immunoblots for lung development transcription factors (as shown in the schematic) in H2087-LCC cells (*left*) and M802T4-LCC cells (*right*), and derivatives of these cells established from spontaneous outbreaks (SO). β-actin was used as loading control.

**Extended Data Fig.2.**
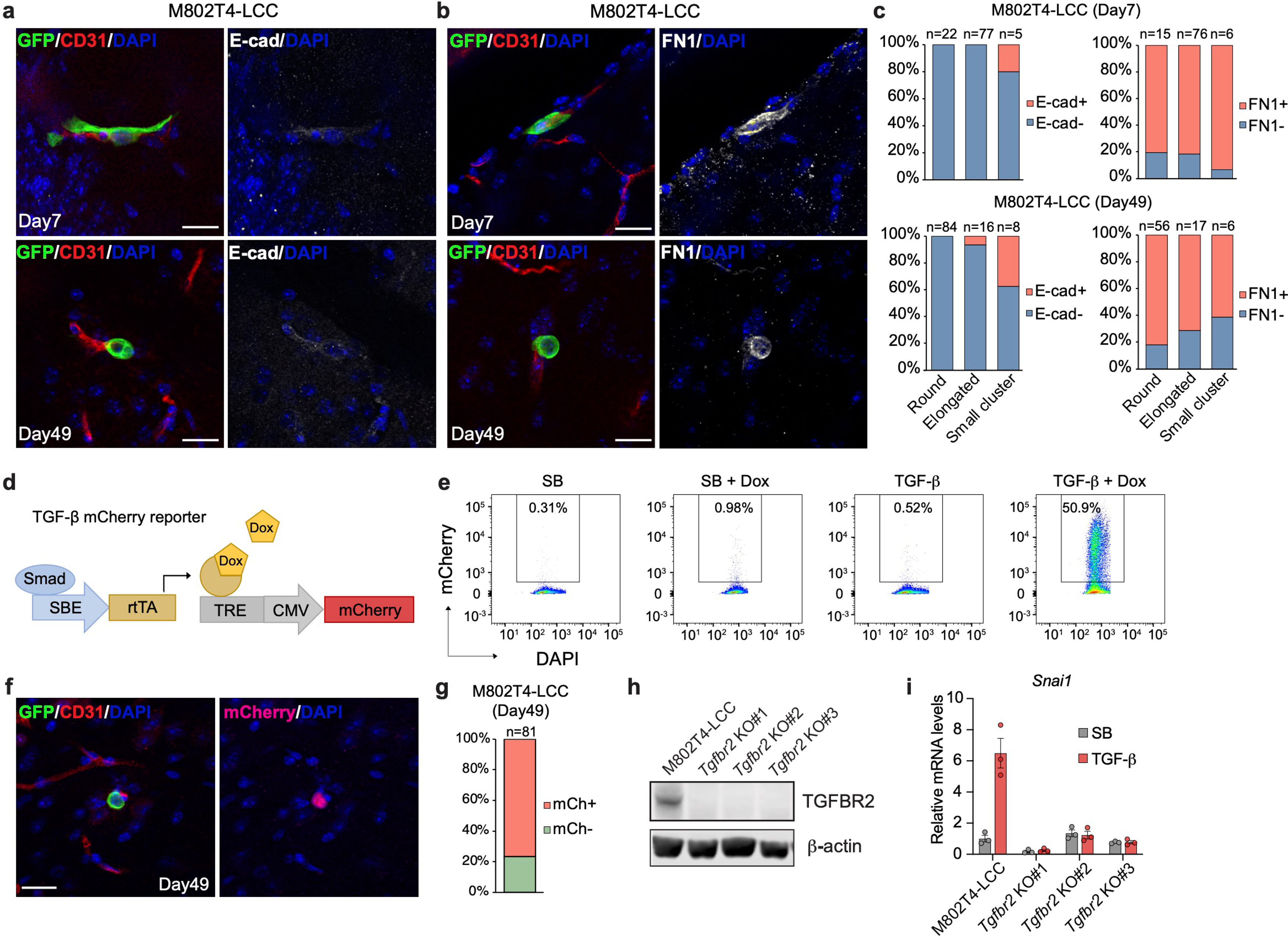
Mesenchymal markers and TGF-β signaling in disseminated LUAD cells. **a** and **b**, Representative IF images of M802T4-LCC cells in brain parenchyma at the indicated time point after intracardiac injection into B6-albino mice. GFP^+^ cancer cells (green), CD31^+^ blood capillaries (red), E-cadherin or FN1 (white), and DAPI (blue). Scale bar: 20 μm. **c**, Quantification of E-cadherin and FN1 IF in brain metastatic 802T4-LCC cells 1 week and 7 weeks after intracardiac inoculation. n= 5 mice per group. **d**, Schematic representation of a doxycycline-inducible TGF-β/SMAD signaling reporter construct driving expression of mCherry. *SBE*, SMAD binding elements; *rtTA,* reverse tetracycline-controlled transactivator; *TRE*; Tet Response Element; *CMV*, cytomegalovirus promoter; *Dox,* doxycycline. **e**, Flow cytometry analysis of H2087-LCC cells expressing a TGF-β mCherry reporter under the indicated conditions. **f**, Representative IF images of M802T4-LCC cells expressing the TGF-β reporter in the brain of B6-albino mice 7 weeks after intracardiac injection. GFP^+^ cancer cells (green), mCherry (magenta), CD31^+^ blood capillaries (red), and DAPI (blue). Scale bar: 20 μm. **g**, Quantification of TGF-β mCherry reporter expression in M802T4-LCC cells (n=81) in the brain of B6-albino mice 7 weeks after intracardiac injection. **h**, Western immunoblot of TGFBR2 expression in wild-type and *Tgfbr2* KO M802T4-LCC cells. β-actin was used as loading control. **i**, qRT-PCR analysis of *Snail* expression in wild-type and *Tgfbr2* KO M802T4-LCC cells upon incubation with SB505124 (SB) or TGF-β for 2 h. Data are mean ± SEM; n=3.

**Extended Data Fig.3.**
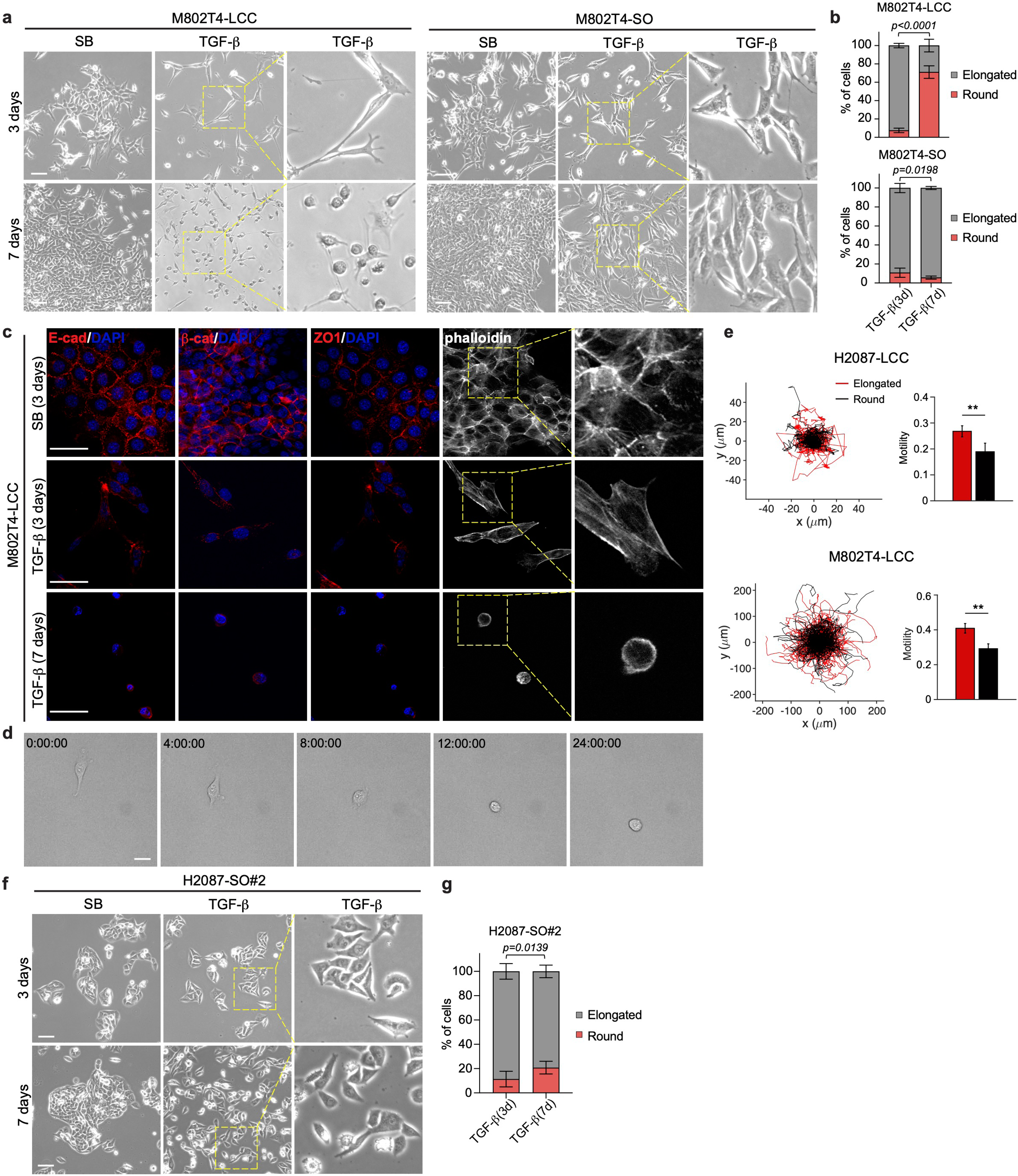
**An EMT state lacking actin stress fibers**. **a**, Representative bright-field images of M802T4-LCC and M802T4-SO cells cultures after incubation with TGF-β for 3 days and 7 days. Scale bar: 200 μm. **b**, Quantification of elongated and spheroidal cells in A. Data are mean ± SD; two-sided unpaired *t*-test. **c**, Representative IF images for E-cadherin, β-catenin, and ZO-1, and of phalloidin staining in M802T4-LCC cells incubated with TGF-β for 3 days or 7 days. Scale bar: 20 μm. **d**, Representative still frames from time-lapse imaging of H2087-LCC cells incubated with TGF-β for 5 days and imaged for 24 h in the presence of TGF-β. Scale bar: 50 μm. **e**, Spider plots of migration tracks and quantification of motility in elongated LCC cells (n=139 and n=319 for H2087 and M802T4 respectively) after 3 days of incubation with TGF-β and spheroidal LCC cells (n=64 and n=282 for H2087 and M802T4 respectively) after 7 days of incubation with TGF-β. **f**, Representative bright-field images of H2087-SO#2 cells upon incubation with TGF-β for 3 days and 7 days. Scale bar: 200 μm. **g**, Quantification of elongated and spheroidal H2087-SO#2 cells upon TGF-β treatment for 3 days or 7 days. Data are mean ± SD.; two-sided unpaired *t*-test.

**Extended Data Fig.4.**
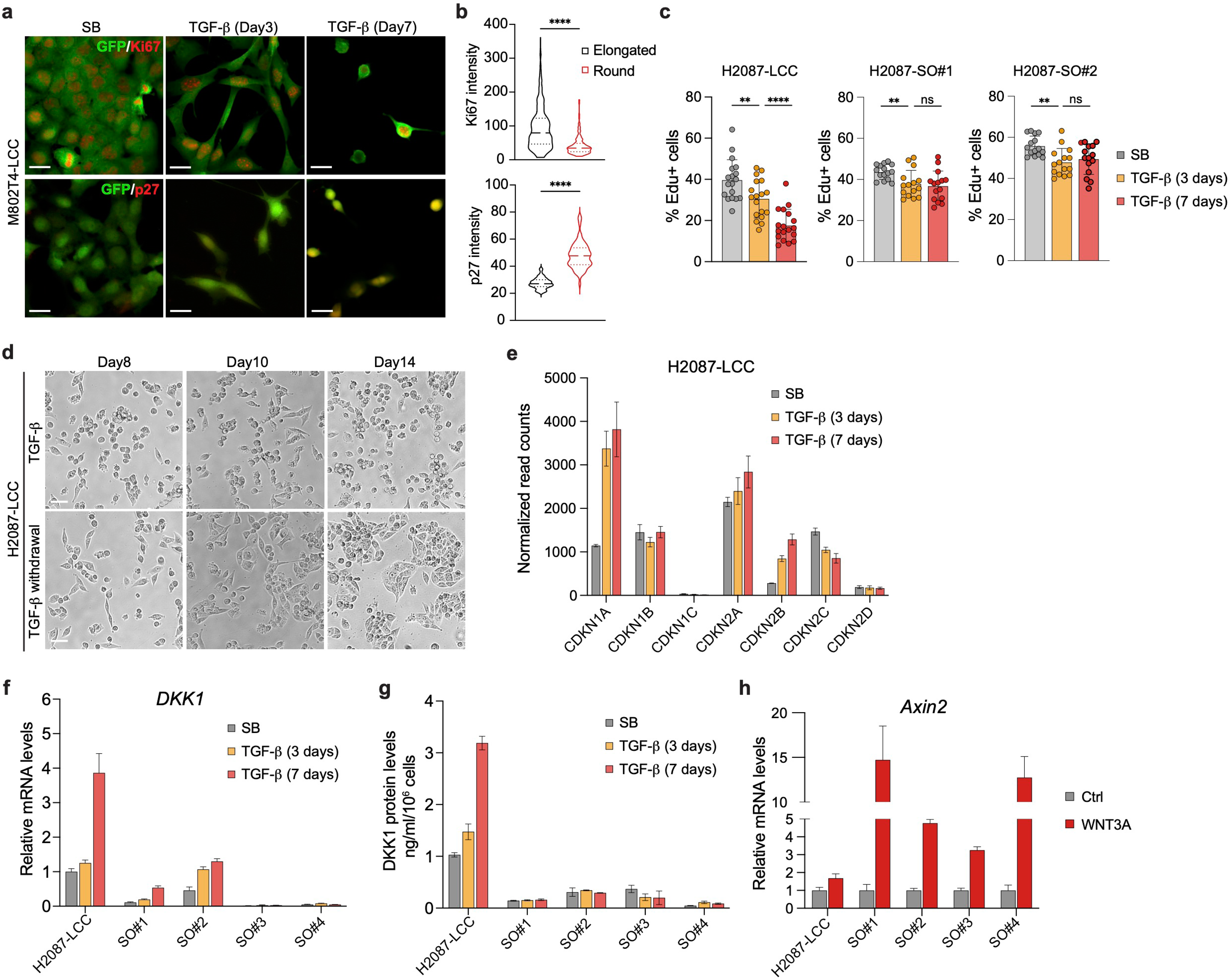
Quiescence-associated EMT responses to TGF-β. a, Representative IF images of M802T4-LCC cells after incubation with TGF-β for 3 days or 7 days. GFP^+^ cancer cells (green), Ki67 (red) or p27KIP1 (red). Scale bar: 20 μm. **b**, Quantification of Ki67 and p27 IF intensity in elongated M802T4-LCC cells after 3 days of incubation with TGF-β and spheroidal M802T4-LCC cells after 7 days of incubation with TGF-β. n=513 (elongated) and n=131 (round) for Ki67 staining, n=121 (elongated) and n=63 (round) for p27 staining. Two-sided unpaired *t*-test. **c**, Quantification of Edu+ cells in H2087-LCC and SO cells upon TGF-β treatment for 3 days and 7 days. Results from 15-18 images for each condition. Data are mean ± SD.; two-sided unpaired *t*-test. **d**, Representative bright-field images of H2087-LCC cell cultures pretreated with TGF-β for 7 days and then switched to media containing TGF-β or SB for the indicated time periods. Scale bar: 200 μm. **e**, Normalized read counts of indicated gene from RNA-seq results of H2087-LCC cells treated with TGF-β for 3 days and 7 days. **f**, qRT-PCR analysis of *DKK1* mRNA levels in H2087-LCC and SO cells treated with TGF-β for 3 days and 7 days. Data are mean ± SEM; n=3. **g**. DKK1 protein levels measured by ELISA in H2087-LCC and SO cells treated with TGF-β for 3 days and 7 days. Data are mean ± SD; n=2. **h**, qRT-PCR analysis of *Axin2* mRNA levels in H2087-LCC and SO cells treated with recombinant human WNT3A for 2 hours. Data are mean ± SEM; n=3.

**Extended Data Fig.5.**
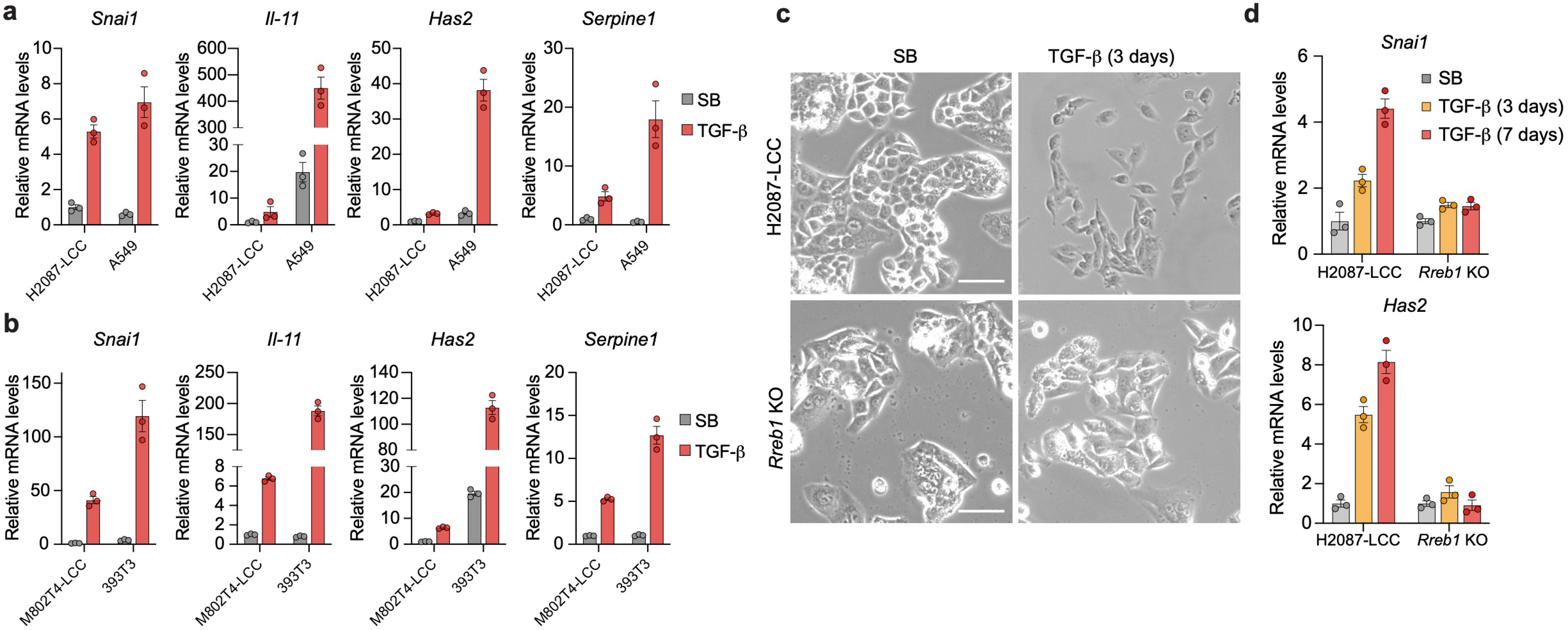
Distinct fibrogenic-EMT responses in different LUAD progenitor states. **a** and **b**, qRT-PCR analysis of *Snai1* EMT-TF and three representative EMT-associated fibrogenic factors^35^ in the indicated cell lines upon TGF-β treatment for 2 h. Data are mean ± SEM; n=3. **c**, Representative bright-field images of wild-type and *Rreb1* KO H2087-LCC cells incubated with TGF-β for 3 days. Scale bar: 200 μm. **d**, qRT-PCR analysis of *Snai1* and *Has2* expression in wild-type and *Rreb1* KO H2087-LCC cells incubated with TGF-β for 3 days or 7 days. Data are mean ± SEM; n=3.

**Extended Data Fig.6.**
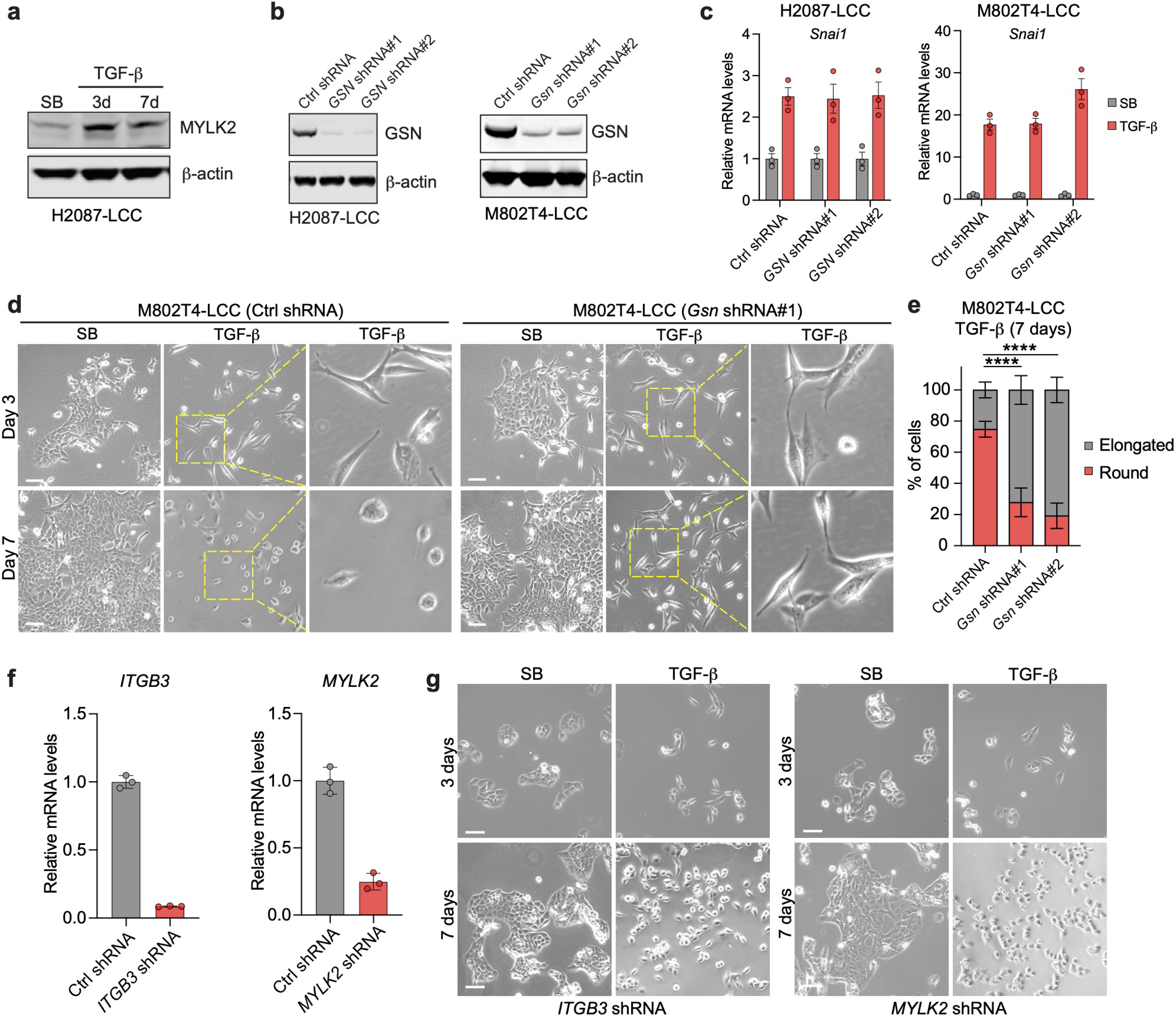
A gelsolin modified EMT. **a**, Western immunoblot analysis MYLK2 levels in H2087-LCC treated with TGF-β for the indicated time periods. β-actin was used as loading control. **b**, Western immunoblot analysis of GSN levels in H2087-LCC and M802T4-LCC cells expressing control shRNA or different shRNAs against *GSN*. β-actin was used as loading control. **c**, qRT-PCR analysis of *Snai1* mRNA levels in H2087-LCC and M802T4-LCC cells expressing the indicated shRNAs incubated with TGF-β for 2h. Data are mean ± SEM; n=3. **d**, Representative bright-field images of M802T4-LCC cells expressing control or *GSN* shRNAs and incubated with TGF-β for 3 days or 7 days. Scale bar: 200 μm. **e**, Quantification of elongated and spheroidal cells in the indicated cell lines upon incubation with TGF-β for 7 days. Data are mean ± SD.; results from 10 images per condition, two-sided unpaired *t*-test. **f**, qRT-PCR analysis of *ITGB3* and *MYLK2* mRNA levels in H2087-LCC cells expressing the indicated shRNAs. Data are mean ± SEM; n=3. **g**, Representative bright field images of H2087-LCC cells expressing the indicated shRNAs and incubated with TGF-β for 3 days or 7 days. Scale bar: 200 μm.

**Extended Data Fig.7.**
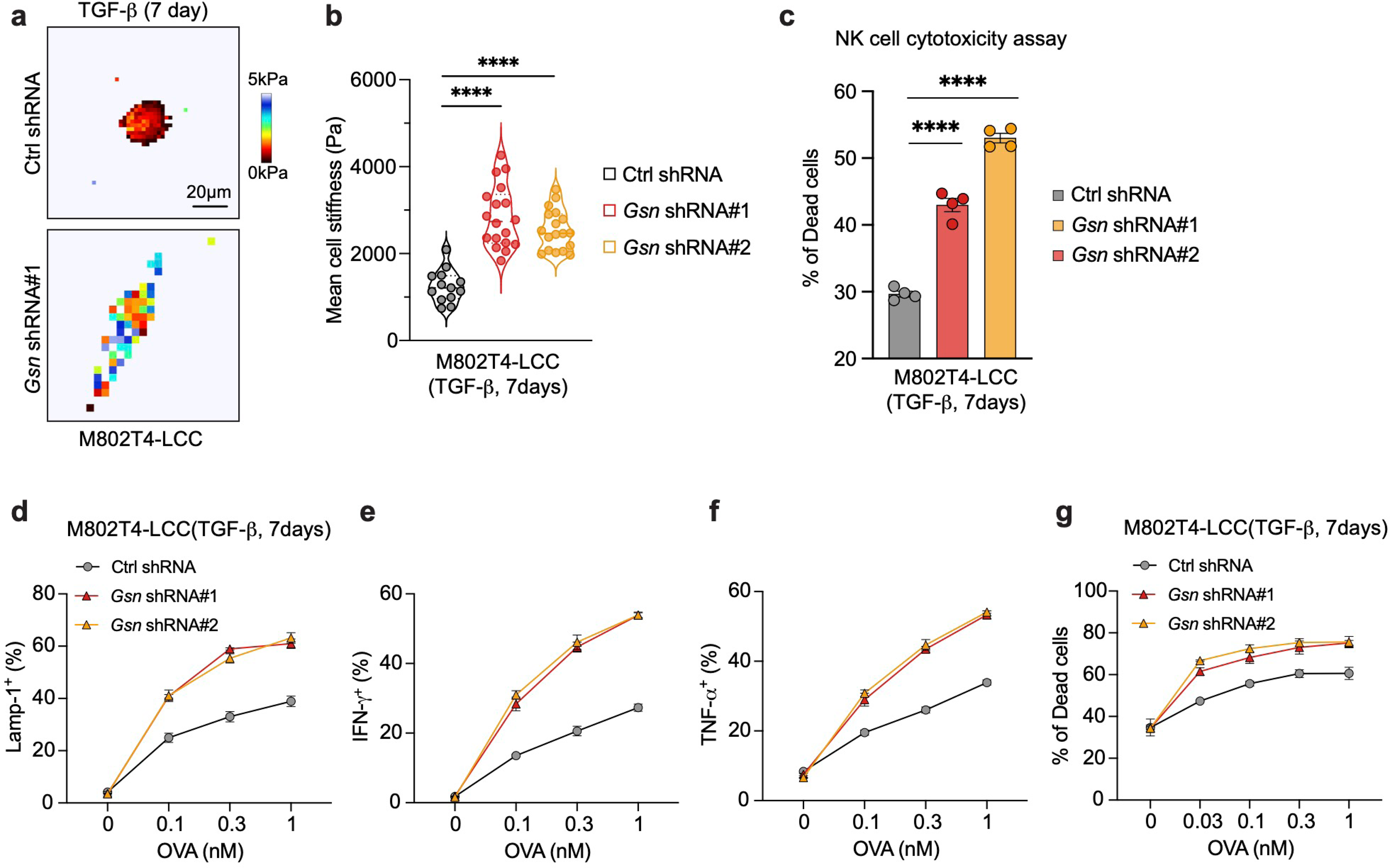
Gelsolin protects LUAD progenitors from mechanosurveillance. **a**, Force maps of representative M802T4-LCC cells expressing the indicated shRNAs and incubated with TGF-β for 7 days. **b**, Mean cell stiffness of M802T4-LCC cells expressing the indicated shRNA and incubated with TGF-β for 7 days. n=13-18 cell per condition; two-sided unpaired *t*-test. **c**, cytotoxic effect of mouse NK cytotoxicity on M802T4-LCC cells expressing the indicated shRNAs and incubated with TGF-β for 7 days. Data are mean ± SEM; n=4; two-sided unpaired *t*-test. **d**, M802T4-LCC cells expressing the indicated shRNA were incubated with TGF-β for 7 days and loaded with OVA peptide. The cancer cells were then co-incubated with OT1 CTLs for 5 h. CTL degranulation was determined by surface exposure of Lamp1. **e** and **f**, Production of IFN-γ and TNF-α, measured by intracellular immunostaining of CTLs from the experiments in D. **g**, cytotoxic effect of OT1 CTLs on M802T4-LCC cells expressing the indicated shRNA and incubated with TGF-β for 7 days. Data are mean ± SD; n=3.

## Supplemental information

**Table 1: GSEA of Kegg pathway dataset in H2087-LCC cells treated with TGF-β for 7 days versus 3 days, related to Figure 2**.

**Video 1: Morphological change of H2087-LCC cells in the presence of TGF-β, related to Figure 2**. A time-lapse movie of H2087-LCC cell pretreated with TGF-β for 5 days and imaged in the presence of TGF-β for 36 hours. Time in HH:MM is indicated in the upper left corner. Scale bar: 50 μm.

**Video 2: NK cell-mediated killing of TGF-β-treated H2087-LCC cell, related to Figure 4**. A time-lapse movie of NK cells attacking a round and an elongated H2087-LCC cells pretreated with TGF-β for 5 days in the presence of Propidium Iodide (PI). Target cell death is associated with structural collapse and PI influx (red fluorescence). Time in HH:MM is indicated in the upper left corner. Scale bar: 50 μm.

